# Combinatorial Wnt signaling determines wing margin color patterns of the swallowtail butterfly ground plan

**DOI:** 10.1101/2024.05.13.593716

**Authors:** Anyi Mazo-Vargas, Alan Liang, Brian Liang, Jeanne M.C. McDonald, Arnaud Martin, Robert D. Reed

**Affiliations:** Duke University Department of Biology, Duke University, Durham, NC 27708, USA; Department of Biological Sciences, The George Washington University, Washington, DC 20052, USA; Department of Ecology and Evolutionary Biology, Cornell University, Ithaca, NY 14853, USA

## Abstract

The intricate wing patterns of butterflies are thought to derive from a morphological ground plan that anchors homology relationships between individual color pattern elements and serves as an archetype for comparative analysis. These patterns undergo modifications that drive the diverse morphologies observed in nature. While brush-footed butterflies (Nymphalidae) have been well studied, assigning homologies with other lepidopteran families remains challenging due to substantial divergence. Here, we focus on swallowtails (Papilionidae), an early-diverging butterfly lineage known for its outstanding diversity in wing shapes and patterns but lacking a developmental framework. Through qualitative and phylogenetic analyses, CRISPR perturbation assays, and *in situ* expression experiments, we investigate homologies between papilionid butterflies, offering phylogenetic and molecular characterization of the Papilionidae wing ground plan. Our results highlight the roles of *WntA* and *Wnt6* in patterning papilionid signature wing elements, such as the glauca and the Submarginal spots. Notably, the nymphalids’ distinct Central Symmetry System is either reduced or absent in the family, with marginal systems expanding proximally. Our data illuminate a highly adaptable patterning system driven by Wnt signaling pathways in developing butterfly wings.

## Introduction

Butterfly wing patterns provide an extraordinary example of rapid diversification of a complex patterning system. These intricate designs are shaped by alterations in the form, color, position, and size of a core set of discrete pattern elements that can be homologized between distant species (Nijhout, 1991), akin to the vertebrate limb ground plan (Hinchliffe, 2002). Early attempts at understanding these patterns were pioneered by Schwanwitsch and Süffert, who proposed ground plan models based on morphological examinations, providing a valuable framework for discerning homologies within butterfly and moth wings (Schwanwitsch, 1924; Schwanwitsch, 1926; Schwanwitsch, 1956; Süffert, 1927). Among these, the Nymphalid Ground Plan (NGP, Fig. 1A) stands out as the most extensively studied prototype within Lepidoptera, offering concrete hypotheses essential for exploring developmental and evolutionary questions, such as trait homology, gene function evolution, evolutionary novelty, and modularity (Monteiro et al., 2006; Suzuki, 2013; Martin and Reed, 2010; Oliver et al., 2012; Martin and Reed, 2014; Mazo-Vargas et al., 2017).

**Figure 1.**
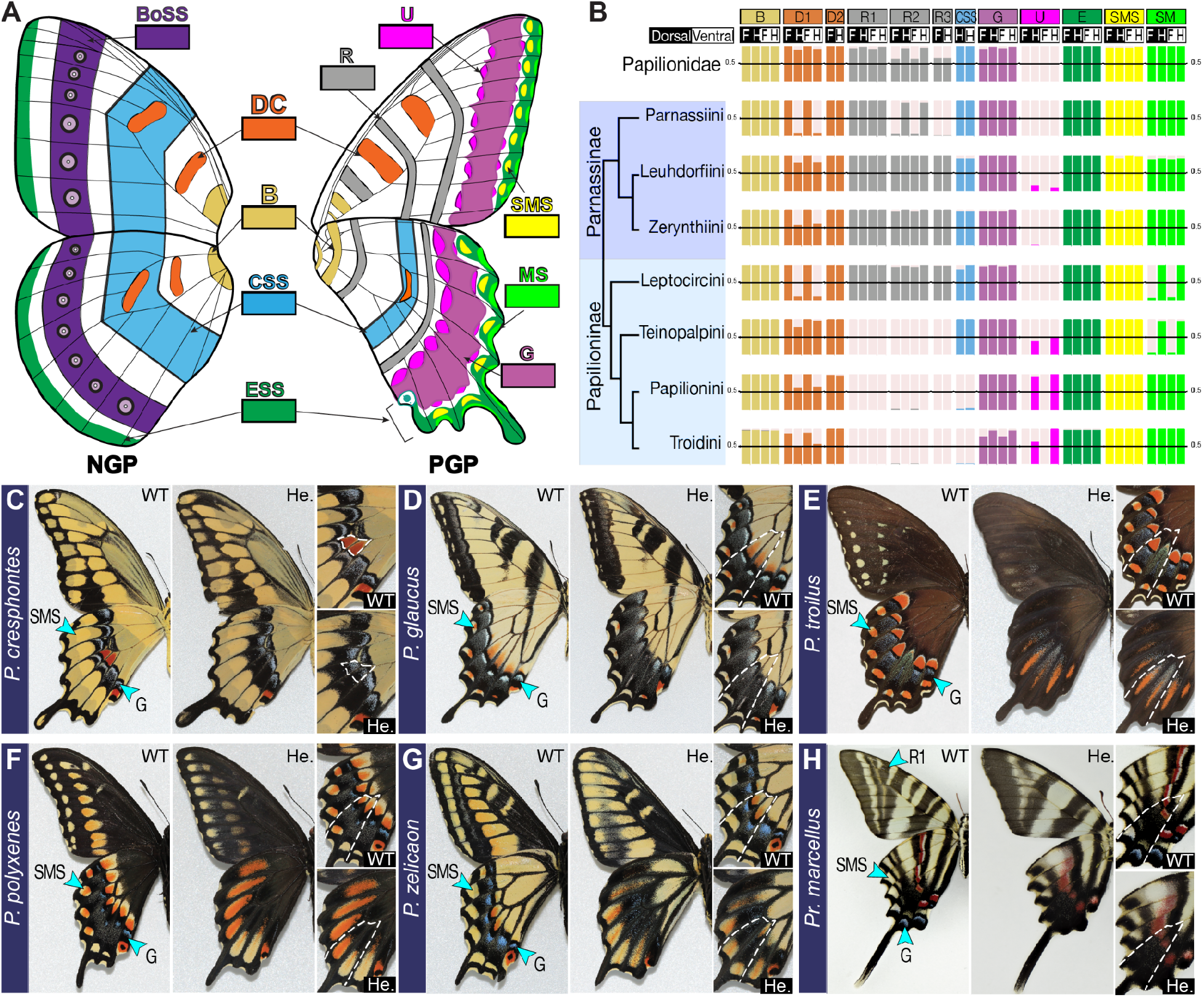
Proposed wing pattern ground plan homologies in butterflies and the effect of heparin perturbations in Papilionidae color patterns. (A) Nymphalid (NGP) and updated Papilionid Ground (PGP) elements derived from (Nijhout, 1991; Schwanwitsch, 1943; Süffert, 1927) and data provided here. (B) Summary of relative ancestral state likelihoods for papilionid symmetry systems. Wild-type and heparin-induced alteration in ventral wing patterns of males of (C) *P. cresphontes*, (D) *P. glaucus*, (E) *P. troilus*, (F) *P. polyxenes*, (G) *P. zelicaon*, (H) *Pr. marcellus*. Dot lines are vein landmarks for reference. Colors in ‘Lettering’ represent Nymphalid and Papilionid Ground Plan elements. B: *Basalis*, BoSS *Border ocelli Symmetry System*, CSS: *Central Symmetry System*, DC: *Discalia I-II*, ESS: *Externa Symmetry System*, G: *Glauca*, R: *Rubrae I-III*, U: *umbra*, SMS: *Submarginal spots*, MS: Marginal spots. See Appendix I for full definitions of pattern elements.

Swallowtail butterflies are, among others, characterized by their involvement in mimetic rings and sex-linked melanism (Brower, 1958; Thompson et al., 2014; Nishikawa et al., 2015) and represent the earliest diverging branch within the butterfly clade, serving as the sister group to all other butterflies (Allio et al., 2020; Kawahara et al., 2023). While the genetics underlying mimicry in this family has received considerable attention (Brower, 1958; Kunte et al., 2014; Thompson et al., 2014; Nishikawa et al., 2015; Palmer and Kronforst, 2020), deeper investigations into wing pattern homologies remain scarce. The most extensive ground plan proposed for Papilionidae dates back to 1943 (Schwanwitsch, 1943), suggesting a significant variation within this family but also drawing parallels between the NGP and the Papilionid Ground Plan (PGP) (Fig. 1A). In essence, these ground plan hypotheses serve as a classification scheme for wing color elements, organizing them based on the positioning of specific stripes and spots. The elements are symmetric and can stretch across the entire wing length and have been denoted as symmetric systems (Fig. 1A). Progressing from the base to the most distal portion of the wing, the NGP encompasses the Basalis, Discalis, Central, Border Ocelli, and Externa Symmetric System (Fig. 1A). In the PGP, additional elements have been identified that do not share the same properties as those found in nymphalids, such as Rubrae, Glauca, and Umbra (Fig. 1A, B). Thus, these observations predict both homologous and derived color elements between these two distantly related families. However, the need for more developmental data has constrained the scope of comparative research.

The simplicity of the ground plan model and the vast diversity of wing patterns in Lepidoptera have motivated researchers to extend this model to other butterfly and moth families using phenotypic observations (Gardiner and Terblanche, 2010; Suzuki, 2013; Schachat and Brown, 2016; Gawne and Nijhout, 2019; Schachat, 2020). Unfortunately, a lack of developmental data has limited the conclusiveness of this comparative work, in contrast to the expression and knockout studies that have been instrumental in evaluating nymphalid color pattern homologies. The Wnt family of secreted proteins has emerged as a focal point of interest for both nymphalids (Martin and Reed, 2014; Mazo-Vargas et al., 2017; Concha et al., 2019; Banerjee et al., 2023; Hanly et al., 2023) and papilionids (Iijima et al., 2019; VanKuren et al., 2023). Particularly, *WntA* has acquired significant attention due to its remarkable variation in gene expression and patterning role across a wide range of nymphalid species (Mazo-Vargas et al., 2017; Concha et al., 2019; Banerjee et al., 2023). Here, we explore the evolution and diversity of pattern elements within Papilionidae from a ground plan perspective, with the goal of establishing a comprehensive morphological and genetic framework for in-depth homology studies. To achieve this, we employ a combination of phylogenetic and genetic methodologies to investigate and propose an updated PGP. Our approach includes conducting *in situ* hybridization, drug perturbation assays, and CRISPR/Cas9 knockouts targeting *Wnt6*, along with the NGP marker gene, the ligand *WntA*.

## Results

### Ancestral Reconstructions of Color Patterns in Papilionidae

Most existing ground plan models for various Lepidoptera lineages are derived from phenotypic observations, mainly focusing on positional relationships between pattern elements, wing veins, and natural variations. We employed a similar approach to classify color pattern elements across 74 papilionidae species (Supp. Table 1), and in addition, we performed ancestral reconstructions using the binary presence/absence of pattern elements for shape and color in this family (Fig. 1B, Fig. S1-4). Based on Schwanwitsch (1943), ten elements were included: from proximal to distal (Fig. 1A, B), we have the Basilis (B), Discalia 1 and 2 (DC: D1, D2), Rubrae 1-3 (R1, R2, R3), Media or Central symmetry system (CSS), Umbra (U), Glauca (G), Externa (E). Additionally, we noted the presence of two rows of spots located towards the wing margin in many species, which we have designated as Submarginal spots (SMS) and Marginal spots (MS) (Shimajiri and Otaki, 2022). A detailed description of each pattern system is in Appendix I.

Overall, the evolutionary trajectory of color elements within Papilionidae reveals a trend towards pattern loss or complexity reduction, particularly evident in the diminished presence of Rubrae and CSS elements (Fig 1B, S1-4). The Rubrae system, distinctive to papilionids, consists of three central bands. These elements are mainly limited to the Parnassiinae, often appearing as red ocelli-like patterns, as well as Leptocircini (Fig 1B, S3-S4). Similarly, the CSS has also undergone a reduction in prominence and is notably different from that of the NGP. While in nymphalids this region is contained by two separate bands, in papilionids it manifests as a single band, primarily present on the hindwings and highly reduced. The ESS, comprised of marginal stripes known as Externa 1-3 in nymphalids (Nijhout, 1991; Reed et al., 2020), differs in papilionids. Schwanwitsch (1943, 1956) identified Externa 1 (here named Externa), which distinctly borders the distal wing margin and is highly conserved in the family (Fig. 1B, S1-S2). While Externa 2 is absent, Externa 3, known as Glauca, stands alone as a distinct pattern element (Schwanwitsch, 1943), emerging as a prominent feature in papilionids and contributing significantly to pattern diversity (Fig. 1B, Fig. S3-S4). This element presents as a prominent dark band that runs parallel to the wing margin, often featuring an iridescent blue interspace (Fig. 1, Fig. S3-S4).

In our revision of the PGP, we added the Submarginal spots and Marginal spots (Fig. 1A). These rows of spots are located between the Glauca and Externa band and the very distal margin of the wing, respectively (Fig. 1A). Although such patterns have usually been described as merely background areas surrounded by darker patterns (Nijhout 1991), their widespread presence in highly melanic species suggests they are pattern elements (see Appendix I). Both elements are highly prevalent across most lineages (Fig. 1B, S1-S2), with the Submarginal spots presenting outstanding variation in color and size.

### Wing margin morphogen sources set *Papilio* symmetry systems

With our updated PGP, we next aimed to elucidate the molecular mechanisms underlying the formation of distinct pattern features. Given our knowledge of the NGP and evidence linking Wnt signaling to color patterning in mimetic *Papilio* species (Iijima et al., 2019; VanKuren et al., 2023), we utilized morphogen-altering drugs and genetically manipulated Wnt genes to evaluate their role at the ground plan level.

Several major NGP color pattern elements are induced by short- or long-range Wnt signaling (Martin and Reed, 2014; Mazo-Vargas et al., 2017; Hanly et al., 2023). Notably, studies have demonstrated that pharmaceutical treatments can manipulate Wnt signaling, resulting in significant phenotypic alterations of Wnt-associated color patterns (Martin and Reed, 2014; Serfas and Carroll, 2005; Sourakov, 2020). Given the pattern-specific effects of these treatments, drug injections serve as a valuable tool for identifying color patterns likely induced by Wnt signaling. Heparin and dextran sulfate affect the activity of signaling pathways (Perrimon, 2004; Serfas and Carroll, 2005). Heparin mimics heparan sulfate proteoglycans within cells, stabilizing Wnt signaling proteins and promoting gradient formation (Fuerer et al., 2010); hence, increasing concentrations of heparin expand Wnt’s range of effect, mimicking a gain-of-function effect in neighboring cells. Conversely, dextran sulfate produces the opposite effect, the contraction of Wnt-induced color patterns (Serfas and Carroll, 2005; Martin and Reed, 2014).

We sought to test the consequences of heparin injections in six Papilionidae species, members of the Papilionini and Leptocircini tribes (Fig. 1C-H, S5-S10). These treatments presented a range of changes in the mid-wing and margin patterns but not in the basal region of the wings, leaving the Basalis and Discalis systems unaffected. The Glauca, but not the Externa, was drastically altered by the heparin treatments, reducing and inducing different color scales (Fig. 1C-H, S5-S10). Minor effects start with a scattering of blue scales and a proximal movement of the black band, suppressing the red/yellow region described by Schwanwitsch as the Umbrae was evident in the *Papilio* species (Fig. 1D, S5-S10). In more substantial cases, there is a drastic reduction of the melanic pattern, allowing the extension of the color pattern from the distal neighbor spots (Fig. 1F-G). These species-specific results were also associated with relative injection time and concentration. Hence, injections close to the moment of pupation and higher amounts displayed stronger color pattern perturbations, spreading melanin across most of the wing. Surprisingly, in *P. polyxenes*, the heparin treatment revealed sex-dependent effects (Fig. S8). This species is sexually dimorphic dorsally but not ventrally. However, in the ventral side of females, the blue and black patterns seemed dominant over the yellow and red elements, contrasting with male effects, suggesting somewhat different pathways specifying similar wing designs between females and males.

*Protographium marcellus*, also known as the zebra swallowtail, offers an intriguing case study within the tribe Leptocircini (Fig. 1B, H, S10). Unlike the *Papilio* species, this tribe displays a diverse array of elements that have not undergone significant reduction. The hindwings of *Pr. marcellus* are predicted to retain remnants of the CSS bordered by Rubrae elements, a feature induced by the gene *WntA* in nymphalids. Interestingly, injections of heparin affected not only the Glauca and Submarginal spots but also the CSS, the forewing and hindwing Rubrae, and the Marginal spots (Fig. 1H, S10).

The Submarginal spots were consistently affected in all species studied. Traditionally viewed as background coloration between the Glauca and Externa, these spots exhibited unexpected behavior under the influence of heparin. In butterflies experiencing strong heparin effects, we observed a surprising expansion of these spots towards the proximal region of the wing, often dominating the wing space and displacing the Glauca. *P. cresphontes*, however, which already feature enlarged spots, did not exhibit any extension beyond their typical boundary (Fig. 1C). In *Papilio* species, color pattern movement and replacement were observed from the wing margin towards the proximal region, suggesting the extension of morphogen gradients along the wing margin (Fig. 1C-G, S5-S9). Conversely, in *Pr. marcellus*, the movement of colors was less clear, possibly due to morphogen sources located in both the middle and margin of the wing (Fig. S10).

### *WntA*, a key ground plan marker, patterns the Glauca in *Papilio* butterflies

We have identified *WntA* as the ground plan gene marker for nymphalid butterflies. In caterpillar imaginal discs it presents up to three parallel expression domains (Martin and Reed, 2014; Mazo-Vargas et al., 2017), which are further refined early in pupal development (Hanly et al., 2023). Based on our findings from heparin injections and previous work in *P. polytes* and *P. alphenor* (Iijima et al., 2019; VanKuren et al., 2023), we hypothesized that *WntA* might also play a role in wing patterning in Papilionidae. To investigate this hypothesis, we examined the spatial expression patterns of *WntA* in caterpillar wing imaginal discs and pupal wings and utilized CRISPR/Cas9 to knock out its function in two species: *P. zelicaon* (Fig. 2A-D, S12) and *P. polyxenes* (Fig. 2E-I, S13). Our results unequivocally demonstrate that *WntA* is indispensable for patterning the Glauca element in both species. Glauca is one of the most prominent features in Papilionidae wings that often exhibits iridescent blue scales (Fig. 2D, I, S1, S4). *WntA* patterning initiates in the caterpillars, where mRNA is notably observed as a relatively thick band near the margin of the imaginal disc of both *P. zelicaon* (Fig. 2A) and *P. polyxenes* (Fig. 2E), with expression on both the proper wing and the peripheral tissue. Very early in pupal development *WntA* signaling activates and patterns are fine-tuned, forecasting details observed on adult wings. In the absence of *WntA* function due to CRISPR knockouts, the Glauca is lost, leading to an expansion of the yellow and/or red background coloration. The yellow background is evident in the forewings of *P. zelicaon* (Fig. 2B, C, S12) and on the dorsal side of male *P. polyxenes*, spanning both forewings and hindwings (Fig. 2E, S13). Conversely, the red background pattern is apparent on the ventral hindwings of both species (Fig. 2D-I, S12-S13).

**Figure 2.**
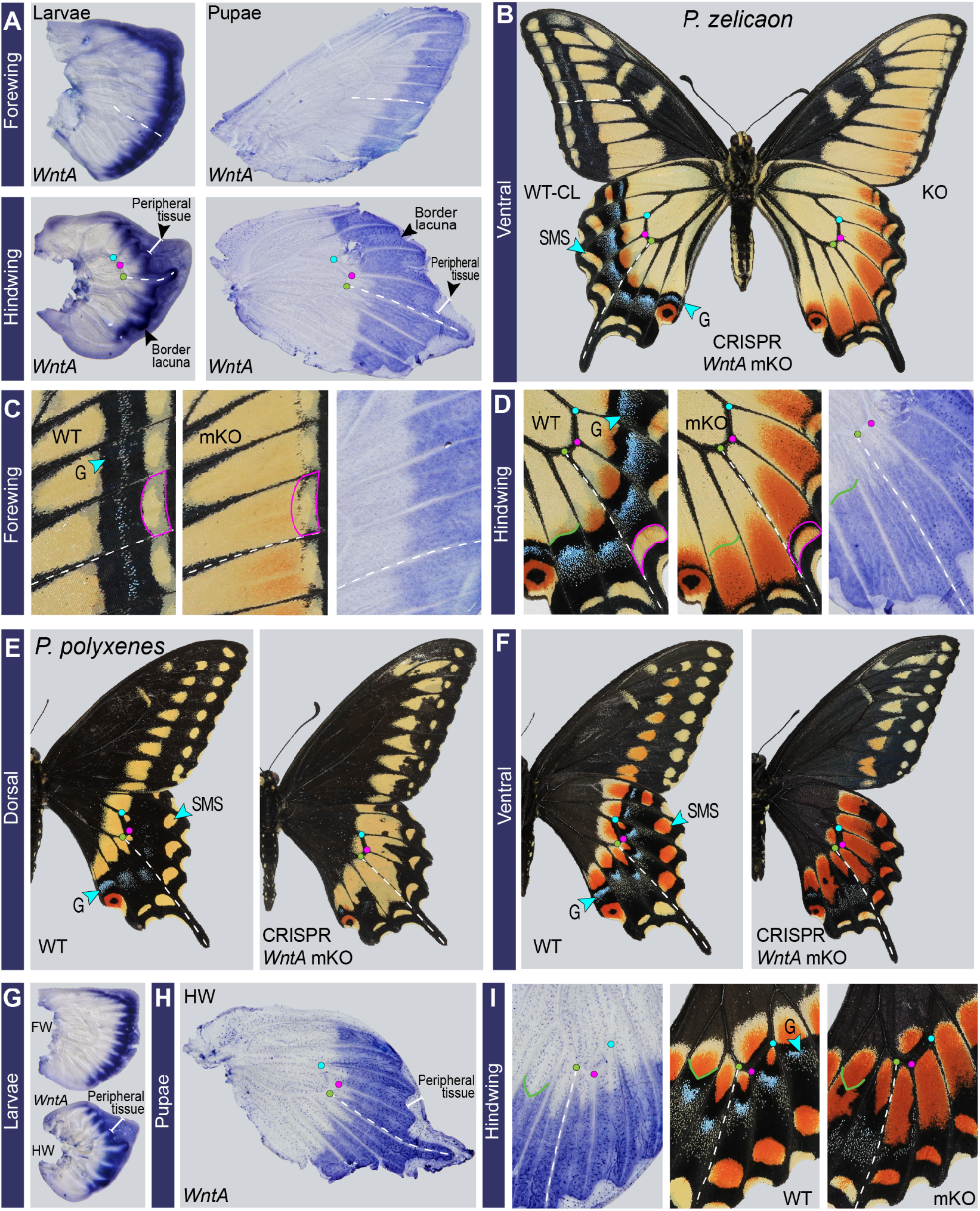
*WntA* specifies the Glauca color pattern system in *Papilio*. (A) *In situ* hybridization of *WntA* in mid-last instar imaginal discs and ∼20% pupal development of *P. zelicaon* in forewing (FW) and hindwing (HW). (B) Ventral adult mosaic knockout (mKO) phenotype after *WntA* CRISPR/Cas9. This individual shows complete left-right differences with a mutant side (KO) and a side devoid of phenotypic effects (WT-CL: wild-type contralateral). (C-D) Close-up of *P. zelicaon* adult phenotypes and pupae *in situ* from panels A and B. Magenta lines highlight differences in the Submarginal spots. *WntA* RNA expression on the pupal wing reflects the shape of the proximal margin of G (Glauca), marked by the green line. (E-F) Dorsal and ventral view of *P. polyxenes* wild type and mKO phenotypes. (G) Larvae and (H) hindwing pupae expression of *WntA*. (I) Close-up ventral hindwing of *P. polyxenes*. Colored dots are landmarks of vein intersections. The dotted white line marks vein M3 in all images.

Interestingly, the loss-of-function results obtained through CRISPR mirror the observations following gain-of-function treatment with higher concentrations of heparin, which poses a contradiction in the outcomes. However, this phenomenon could potentially be elucidated by *WntA* suppressing an unidentified neighboring morphogen source, which disperses after either gene activity removal or when heparin concentrations reach a certain threshold. These findings may suggest the presence of at least two distinct signaling sources in *Papilio*. Intriguingly, the knockouts revealed no discernible sex differences in the ventral hindwings of *P. polyxenes* (Fig. S13), contrary to observations made during heparin treatments, indicating the involvement of a different signaling molecule in female-specific patterning. Lastly, while no effects were detected in the basal portion of the wing in *WntA* knockouts, a slight reduction in the size and shape of the Submarginal spots was observed (Fig. 2C-D, I), as discussed below.

### *WntA* and *Wnt6* shape the Submarginal spots in *Papilio*

Heparin injections and *WntA* knockouts suggest the presence of a previously unrecognized wing pattern system between the Glauca and the Externa bands. Typically, in butterflies, the wing ground plan elements are described in terms of pattern elements over a ‘plain’ background (Nijhout, 1991; Otaki, 2008), where the color patterns are pigmented designs determined by concentration levels of various inductive molecules (Nijhout, 1991; Serfas and Carroll, 2005; Otaki, 2011; Martin and Reed, 2014)

After knocking out *WntA*, we noticed a row of round or semi-rectangular Submarginal spots running along the margins of *P. zelicaon* forewings (Fig. 2B-C), but showing yellow coloration similar to the background. Close examination of both forewings and hindwings in *P. zelicaon* and *P. polyxenes* show asymmetric size and shapes between contralateral views of the same individual, ruling out any type of individual variation (Fig. 2B-D, I). Moreover, these spots appeared slightly compressed distally, contrasting with their proximal extension observed in the heparin treatment (Fig. 1F-G). To explore the role of Wnt signaling further, we administered dextran sulfate injections, which antagonize Wnt signaling. We observed phenotypic alterations in *P. zelicaon* (Fig. 3B, D), notably a distal movement of the Glauca coupled with a reduction, sometimes severe, of the Submarginal spots. Thus, our results collectively suggest a non-*WntA* ligand could underlie the specification of the Submarginal spots – i.e., drugs that modulate *Wnt* signaling affect the spots, but *WntA* knockouts do not cause loss of the spots, only subtle shape and size changes (Fig. 2B-C).

**Figure 3.**
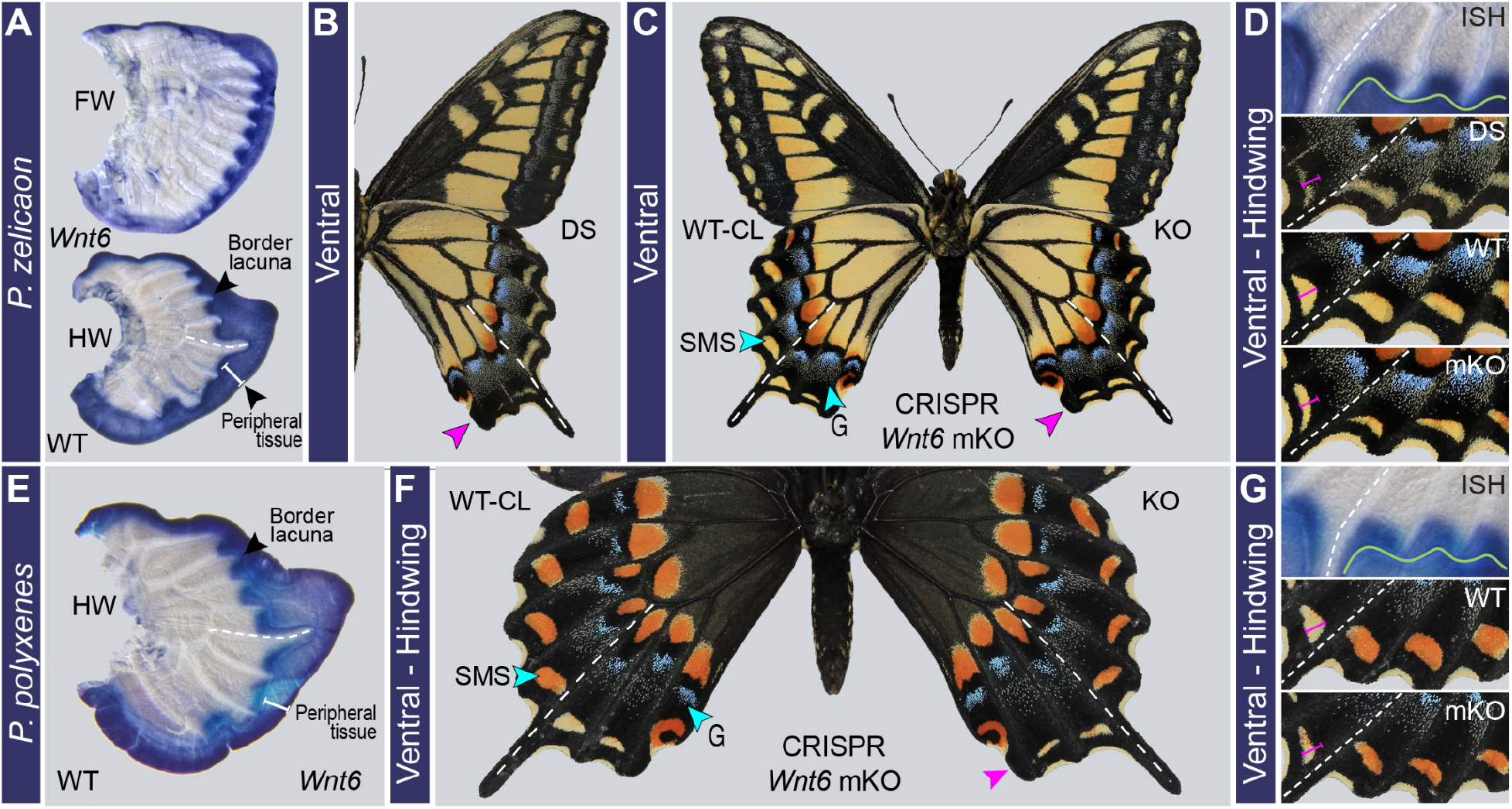
*Wnt6* contributes to patterning Submarginal spots in *Papilio*. (A) In-situ hybridization (ISH) of *Wnt6* on caterpillar fore (FW) and hindwings (HW) imaginal discs of *P. zelicaon*. (B) Dextran sulfate (DS) treatment and (C) *Wnt6* CRISPR/Cas9 knockout of *P. zelicaon*. (D) close-up of the ventral hindwing margin from A, B, and C. (E) ISH of *Wnt6* on caterpillar hindwings (HW) imaginal discs of *P. polyxenes*. (F) *Wnt6* mosaic knockout in *P. polyxenes* ventral hindwing. (G) Close-up of wing margin from E and F. Magenta arrows and lines highlight the reduction in the size of the Submarginal spots (SMS). The dotted white line is the landmark M3 vein, and the green line corresponds to the Border lacuna (future adult wing margin) in the imaginal discs.

During wing development, dorsal and ventral epithelial cell layers fuse in the last larval instar, leaving a system of trachea cavities (Nijhout, 1991). This creates the *border lacuna* running parallel to the imaginal disc margin that shapes the future adult wing (Fig. 2A - 3A). Between these two boundaries is found the *peripheral tissue*, which undergoes apoptosis during the pupal stage. The marginal color patterns observed in the adult butterflies are then proximal to the *border lacuna*. wg/Wnt1 and the transcription factor cut mark the peripheral tissue in Lepidoptera (Macdonald et al., 2010; Martin and Reed, 2010) (Fig. 4A), with several *Wnt* genes known to be expressed along this region in nymphalid butterflies (Martin and Reed, 2014; Banerjee et al., 2023). RNAi knockdown experiments of *Wnt1* and *Wnt6* demonstrated subtle effects on the Externa and the Submarginal spots, respectively (Iijima et al., 2019). We examined the expression of one of these genes, *Wnt6*, and found that in both *P. zelicaon* and *P. polyxenes*, it is transcribed along the border lacuna as discrete spots, with an elongated and semi-rectangular shape, correlate with the adult Submarginal spots pattern (Fig. 3A, D-E, G). Next, we used CRISPR/Cas9 deletion to test the function of *Wnt6* in color patterning. The knockouts caused a drastic reduction of the Submarginal spots in both *P. zelicaon* (Fig. 3C-D, S14) and *P. polyxenes* (Fig. 3F-G, S15). The mutant phenotypes consisted of changes in shape and size of the spots, but not color, similar to the *WntA* effect. Our data highlight these Submarginal spots in *Papilio* as true, dominant pattern elements instead of ‘plain’ background, where *WntA* and *Wnt6* are positive regulators of element size and shape.

**Figure 4.**
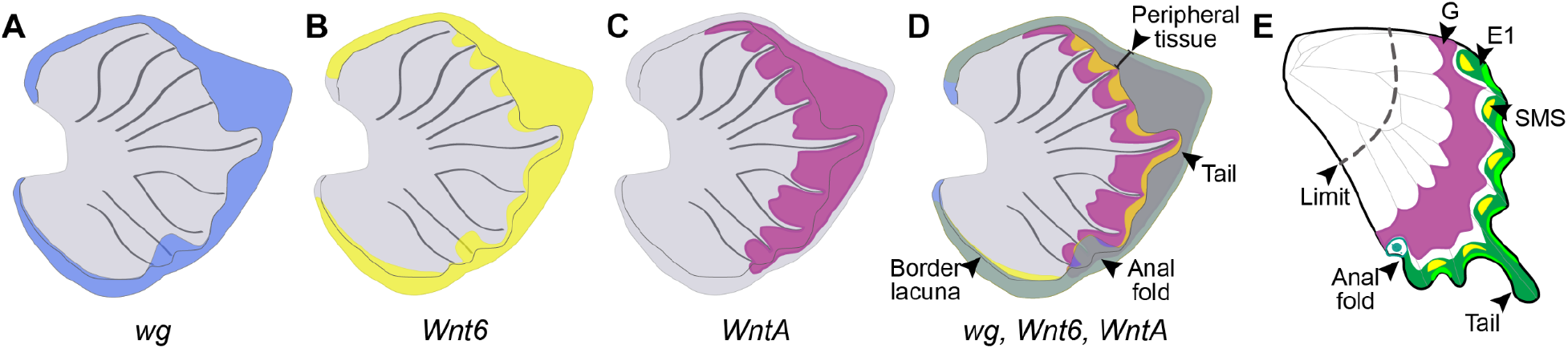
Combinatorial Wnt signaling induces margin color pattern elements in *Papilio*. Schematic representation of hindwing imaginal disc from the last larval instar illustrating (A) known *wg/Wnt1* expression predominantly in the peripheral tissue. (B) *Wnt6* mRNA was detected in both the peripheral tissue and the proper wing, which correlated with the Submarginal spots (SMS) position. (C) *WntA* expression was observed in both the peripheral tissue and the proper wing, predicting the Glauca position and shape. (D) The intersection of Wnt ligands shows extensive overlap in the peripheral tissue and gene-specific expression domains, which pattern the margin elements. (E) Glauca and the Submarginal spots elements in the adult wing.

Lastly, one intriguing element is the singular eyespot-like pattern often seen along the margin, at the base of the hindwing tail in numerous papilionid species, with correlated *Wnt6* and *wg/Wnt1* expression during larval development (Fig. 3, 4). This spot seems unaffected by our *WntA* or *Wnt6* knockouts and does not respond to heparin except at very high concentrations, suggesting redundant action of signaling ligands are involved in wing shape but not in inducing the spot.

## Discussion

The ground plan depictions of butterfly and moth wing color patterns are theoretical models that allow us to generate and test hypotheses related to morphological development and evolution. These models were originally derived from observations of phenotypic variation within and between species. Adding new genetic and developmental data now allows us to probe a deeper set of evolutionary questions, including the molecular basis convergence, the evolution of gene function in pattern diversification, and the genetic origins of evolutionary novelties in this megadiverse group of insects. Here, we present a revised model for papilionid wing patterns based on phylogenetics, pharmacologic manipulation of morphogen gradients, *in situ* expression in developing wings, and knockouts of two *Wnt* genes. This model is the first step in gaining insights into the wing patterning in this butterfly family that presents extensive examples of pattern variation, aposematism, and sexual dimorphic wing variation. Furthermore, since papilionids are the oldest diverged clade of butterflies, this work provides an important phylogenetic point of reference for future studies looking at the origin and evolution of the wing pattern ground plan itself.

### Evidence for divergent proximal patterning in Papilionidae

The major ground plan elements described for both nymphalids and papilionids are: Basalis, Discalis (D1, D2), Central Symmetry System (CSS), and the Externa Symmetry System (ESS). Basalis and Discalis were not affected by any of our heparin or CRISPR treatments, suggesting that they may not be induced by Wnt signaling, or at least not the ligands tested here. In the case of Discalis patterns, D1 and D2 are marked by strong *wingless* (*i*.*e*., *Wnt1*) expression in butterflies and moths (Macdonald et al., 2010; Martin and Reed, 2010). Interestingly, in all species we surveyed, heparin-mediated pattern expansions never impacted the proximal region of the wing, as vertical stripes and patterns in about one-third of the wing base remained intact in all experiments (Fig. 1C-H, S5-10, 4E). This division has also been observed in cold-shock experiments, drug perturbations, and naturally occurring phenotypes across Papilionidae (Tyler et al., 1994; Dabrowski and Dobranski, 1996; Umebachi and Osanai, 2003; Scriber et al., 2009; Perlman and Perlman, 2019). In contrast, heparin injections and cold shocks invariably expand the proximal patterns of other lepidopteran lineages, including Nymphalidae, Lycaenidae, and Arctiidae (Iwata et al., 2013; Gawne and Nijhout, 2019; Sourakov, 2020). Thus, the patterning of the proximal region appears to be divergent in Papilionidae, and we speculate that mechanisms other than Wnt signals or heparin-sensitive morphogen drive proximal stripe formation in swallowtail butterflies (Fig. 4E).

### Loss of the Central Symmetry System in *Papilio*

The Central Symmetry System represents one of the largest, most diverse, and significant pattern systems in nymphalid butterflies. Its origins are believed to trace back deeply within Lepidoptera, as similar color patterns are frequently observed across various moth species (Nijhout, 1991; Schachat and Brown, 2016; Gawne and Nijhout, 2019; Schachat, 2020). *WntA* gene expression marks the Central Symmetry System in nymphalids; however, none of our *WntA* expression or perturbation data had any bearing on color patterns in the middle of the *Papilio* wings where the Central Symmetry System would be expected. Thus, we speculate that the Central Symmetry System has been mostly lost in *Papilio*. However, representatives of other tribes, like *Protographium*, are predicted to present a different set of central stripes called the Rubrae (Schwanwitsch, 1943; Schwanwitsch, 1956) in addition to remnants of the CSS elements. Our heparin perturbations favor this hypothesis, but further expression and gene perturbation will be necessary to test and fully understand the evolution and development of the Rubrae and Central Symmetry System in papilionids. Nevertheless, all evidence suggests that the Central Symmetry System is not a major component of the papilionid ground plan, as are the Glauca and the Submarginal spots.

### *Papilio* pattern diversity is largely derived from the Glauca and Submarginal spots

The most striking results from experimental manipulations were in the Glauca. This element was renamed by Schwanwitsch (1956) as Externa III and synonymized by Nijhout (1991) as “parafocal elements” in nymphalids. In the *Papilio* specimens we examined, *WntA* expression extended broadly from the wing margin in both caterpillars and pupae (Fig. 2), and although nymphalids also display an expression domain along the margins, it is less elaborate and spatially restricted (Martin and Reed, 2014; Mazo-Vargas et al., 2017; Hanly et al., 2023). In fact, our knockouts of *WntA* completely removed the Glauca (Fig. 2). In contrast, *WntA* knockouts in nymphalids have much subtler effects on the Externa elements (Mazo-Vargas et al., 2017) – for instance, the Externa III shift distally and are misshapen, which suggests a different developmental underpinning that is influenced by, but not wholly reliant, on *WntA*. This observation suggests that there may have been an ancestral expression pattern for *WntA* associated with the Externa symmetric system, which subsequently evolved in different directions between the two families. However, it remains unclear at this point whether there is direct homology between the *Papilio* Glauca and the nymphalid Externa III elements. With that said, immunostainings of the proteins spalt and Distal-less in the imaginal disc of *P. machaon* (Shirai et al., 2012), a close relative of *P. zelicaon* and *P. polyxenes*, show these proteins are expressed along the wing margin. Both spalt and Distal-less are involved in the formation of Externa I, II, and III in nymphalids, with spalt being necessary for the establishment of the Externa III band (Zhang and Reed, 2016; Reed et al., 2020). Thus, an ancient homology might exist, followed by the subsequent evolution of the developmental network, expanding the expression domains of *WntA* in the margin.

The Glauca manifests as a dark band embellished with blue scales in many papilionid species. However, in certain groups, such as the mimetic *Papilio alphenor*, the Glauca appears black and next to a red pattern in the hindwing. *WntA* RNAi knockdowns reduce both the black and red patterns (VanKuren et al., 2023). Providing more evidence that *WntA* establishes region boundaries, but the pigmentation is controlled by downstream effectors (Mazo-Vargas et al., 2017; Mazo-Vargas et al., 2022; Hanly et al., 2023).

Distal to the Glauca, there is a row of spots that constitutes a dominant pattern system that we term Submarginal spots (Fig. 1A). By examining different species of the family, we noticed the considerable diversity of colors and sizes of these spots, which also are involved in mimicry among different species. For instance, these spots play an essential role in the mimetic species *Papilio polytes*, and knockdown *Wnt6* function with small interfering RNA obtained similar results to our *Wnt6* CRISPR/Cas9 experiments (Fig. 3), i.e., the reduction in size and shape change of the spots (Iijima et al., 2019). We conclude that the interplay between Submarginal spots and the Glauca drives a wide range of the color pattern diversity in papilionid butterflies.

At least three Wnt ligands, including wg/Wnt1, Wnt6, and WntA, exert their influence on the wing margin during development in papilionds (Fig. 4). Among these, wg/Wnt1 demonstrates a highly conserved expression pattern in the peripheral tissue, observed across various species of butterflies and moths (Martin and Reed, 2010). Additionally, *Wnt6* and *Wnt10* expression is also detected in the peripheral tissue in nymphalids (Martin and Reed, 2014; Banerjee et al., 2023). Intriguingly, our findings reveal that not only *Wnt6* but also *WntA* are expressed in the peripheral region in papilionids imaginal discs (Fig. 2A, 2G, 3A, 3E, 4). *WntA* is only expressed in the proper wing for all the nymphalid species studied so far (Martin and Reed, 2014; Mazo-Vargas et al., 2017; Banerjee et al., 2023). This observation leads us to speculate that the ancestral expression domain for the Wnt genes in the wings might reside in the peripheral region. Over evolutionary time, the extension of their expression into the proper wing territory established the spatial cues necessary for generating a diverse arrange of pattern elements.

In *Papilio*, the extension of *WntA* and *Wnt6* into the proper wing, and their interactions regulate the shape and size of the Submarginal spots. Interestingly, mathematical models for *P. dardanus* and *P. polytes* predict that a single gradient originating from the wing margin is sufficient to explain color patterns in these species (Sekimura et al., 2007). Our data largely agree with the central importance of wing margin pattern induction; however, we show clear evidence that gradients of at least two marginal factors – *WntA* and *Wnt6* – interact to induce the *Papilio* color patterns. In species like *Pr. marcellus*, where Central and Rubrae systems are predicted, we observed heparin effects in the middle and margin of the wings. Therefore, we predict the presence of morphogen sources in those wing regions, similar to nymphalids, but without extending beyond a proximal-distal limit.” (Fig. 4D).

We present an updated ground plan for the Papilionidae family (Fig. 1A), offering new insights into developmental mechanisms underlying two major patterning systems, the Glauca and Submarginal spots (Fig. 4). Papiliod wing patterns appears to present a general trend of pattern element loss over phylogenetic time, relying on fewer but more prominent pattern elements to generate their diversity. Notably, both NGP and PGP are predicted to share certain symmetric systems, such as the CSS and ESS, albeit with contrasting refinements. While papilionids predominantly feature ESS elements, nymphalids exhibit more elaborated CSS patterns. It is evident that Wnt signaling plays a predominant role in the evolution of color patterns across various butterfly lineages. Specifically, *WntA* is essential for pre-patterning major elements such as the CSS and ESS during imaginal disc development. Nevertheless, different developmental processes have likely influenced each system’s expansion and diversification (Shirai et al., 2012). Here, we also characterize a novel Submarginal spot pattern system in *Papilio*, in which *Wnt6* and *WntA* signaling play a promoting role. Overall, our findings suggest that much of the color pattern complexity and diversity in *Papilio* can be traced to the interaction between the *WntA* Glauca patterns and the *Wnt6/WntA* Submarginal patterns (Fig. 4), with some occasional ornamentation thanks to Discalis, Rubrae, and/or highly reduced CSS patterns that are not yet well understood in Papilionidae.

## Methods

### Surveying photos and taxon sampling

Photos of the dorsal and ventral aspects of pinned papilionid butterfly specimens were obtained from public sources: https://www.europeana.eu/en/, http://www.globis.insects-online.de/, **Error! Hyperlink reference not valid. Error! Hyperlink reference not valid**. s, http://insecta.pro/ (Table S1). This included a total of 74 species, representing all 31 recognized genera in the family (Allio et al., 2020). Moreover, most of the subgenera of the species-rich genera *Parnassius* (*Parnassius, Lingamus, Driopa*, and *Kriezbergius*), *Graphium* (*Pathysa, Arisbe, Pazala*, and *Graphium*), and *Papilio* (*Heraclides, Alexanoria, Chilasa, Pterourus, Eleppone, Druryia, Sinoprinceps, Papilio, Achillides, Princeps, and Menelaides)* were represented. For polymorphic or sexually dimorphic species such as *Papilio glaucus*, one or more forms may have been included for a total of 81 papilionid specimens (forms are specified on the phylogeny). The nymphalid *Vanessa cardui* (Nymphalidae: Nymphalinae) was included as an outgroup. In total, 164 images (dorsal/ventral aspects of 81 ingroup specimens + 1 outgroup specimen) were used for this analysis. A complete list of species and photograph sources are listed in Table S1.

### Characters

All twelve pattern elements included in this work are described in the Appendix I. They were included as characters: Basilis (B), Discalis I and II (D1-2), Rubrae I, II, and III (R1-3), Media (M) or Central Symmetry System (CSS), Glauca (G), Umbra (U), Externa (E), Submarginal spots (SMS), and Marginal spots (MS). Each of the pattern element characters for the 82 specimens were binarily scored for either presence or absence on the dorsal forewing (DF), dorsal hindwing (DH), ventral forewing (VF), and ventral hindwing (VH). This summed up a total of 48 pattern element characters.

While scoring specimens for various traits, it became apparent that certain pattern elements displayed notable shape and/or color variations. The serially repeated subunits forming R1, G, and E could be roughly categorized into three shapes: band, chevron, and ocellar, with E also classified as an intervenous (IV) stripe. Moreover, R1-3 and G showed discrete variations in the presence or absence of red and blue coloration. The scoring process was repeated to better grasp these pattern elements’ evolutionary trends regarding shape and color, incorporating observed variants for shape and color as additional character states alongside absence. Separate scoring for shape and color were organized into two additional character matrices.

### Ancestral state reconstruction

To understand the evolutionary history of pattern elements, ancestral states for the pattern elements were reconstructed using Maximum Likelihood (ML) criterion and the Markov k-state one-parameter (Mk1) model in Mesquite v. 3.61 (Maddison and Maddison, 2019). The phylogeny of the sampled taxa was based on a recently published phylogeny of 408 species that used seven genetic markers and was constructed using Bayesian methods (Allio et al., 2020). Forty-eight reconstructions were made for the binary presence/absence of pattern elements (all four wing surfaces for all twelve pattern elements), 12 reconstructions were made for shape (all four wing surfaces for pattern elements R1, G, and E), and 14 reconstructions were made for color (all four wing surfaces for pattern elements R1, R1 and G; DF and VF for R3).

### Species sampled for drug and genetic perturbations

We sampled available species distributed throughout the *Papilio* phylogeny, specifically *P. zelicaon, P. polyxenes, P. glaucus, P. troilus*, and *P. cresphontes*, as well as one species of the Leptocircini tribe, *Pr. marcellus* (Fig. 1C-H). Information about the population origin, caterpillar, and adult diet is reported in Table S2. All colonies were kept in a growth chamber at 25-28ºC, 60% relative humidity, and 16/8 h day/night cycle during egg collection or larvae rearing.

### Drug injections

Pupal injections of heparin and dextran sulfate were done as previously described (Martin and Reed, 2014; Sourakov, 2020). Each individual was injected between 4-20 hours after pupation (Table S3), in the basal wing region, taking care to avoid tissue damage. We used hand-pulled glass capillary needles mounted on a micropipette to deliver the drug. Pupae were reared under the conditions described above.

### *In situ* hybridization

We dissected imaginal discs from the last caterpillar instar from *P. zelicaon* in cold PBS. Imaginal discs were stored in TRIzol™ Reagent (Invitrogen™) to extract total RNA using the PureLink™ RNA Mini Kit. cDNA synthesis was then performed using M-MuLV reverse transcriptase (New England Labs). PCR oligos were designed based on conservation between the published *Papilio* genomes (lepbase.org, Table S3). After amplification, pGEM TA cloning was made, with subsequent plasmid verification using Sanger sequencing. The plasmids were linearized and used to create DIG-labeled riboprobe synthesis following the manufacturer’s instructions (Roche). To perform the *in situ* hybridization of the DIG-riboprobes, we dissected imaginal discs and pupae wings to be stained following a previously published protocol (Martin and Reed, 2014).

### Genetic manipulation

We amplified and sequenced *WntA* and *Wnt6* from *P. zelicaon* and *P. polyxenes* genomic DNA and cDNA to aid the design of CRISPR/Cas9 sgRNAs. We verified sequence conservation and used the same sgRNAs for both species (Table S4). Freshly laid eggs were treated with 5% benzalkonium chloride (Sigma-Aldrich) for 90 seconds, then thoroughly washed with water. Eggs were glued to a glass slide. We injected 2 or 4 sgRNAs per gene to knock out the gene function. For each injection, a mix of Synthego synthetized sgRNAs (final 250 ng/ul each) plus recombinant Cas9 protein (0.5ug/ul, PNA Bio) were injected using aluminosilicate glass needles (Sutter). Hatched caterpillars were reared as described above. G0 butterflies were inspected for mutant phenotypes; in this case, the presence of mutant clones in the wings. For adults showing distorted phenotypes were pinned and photographed using a Canon EOS 60D, with a lens Canon EF 100mm f/2.8L USM Macro Lens.

## Acknowledgments

We thank Anna Ren for helping with butterfly husbandry and facilitating the collection of immatures of zebra swallowtails. This work was supported by the following awards: DGE-1650441, DBI-2109536, and IOS-1656514.

