## supplemental file for "Combinatorial Wnt signaling determines wing margin color patterns of the swallowtail butterfly ground plan"

### Supplementary Materials

#### Appendix I
#### Definitions of pattern element nomenclature

1. **Basilis (B):** The proximal-most band near the base of the wing.
2. **Discalis I (D1):** A dark spot or bar pattern on the discal cell crossvein, often referred to as the "discal spot".
3. **Discalis II (D2):** A dark spot or bar pattern near the center of the discal cell, delineated anteriorly-posteriorly by the veins of the discal cell. D2 is evidently absent on the hindwing of papilionids.
4. **Rubra I (R1):** Part of a system of three central bands that alternate in position with the Discalis elements and are often longitudinally split by red. R1 is the distal-most of the three rubral bands, positioned immediately distal of the discal cell crossvein. However, it is often dislocated on the forewing such that the portion of the band posterior of the discal cell is located more proximally and lines up with D1.
5. **Rubra II (R2):** The middle of the three rubral bands. On the forewing, it passes through the center of the discal cell and is positioned between D1 and D2 if they are present. On the hindwing, it is usually located relatively proximally and often lines up with the forewing R3. Like R1, R2 is sometimes dislocated on the forewing such that the posterior portion of the band is located more proximally and lines up with D2.
6. **Rubra III (R3):** The proximal-most of the three rubral bands. On the forewing, it is positioned between D2 and B if they are present. It is absent on the hindwing of papilionids.
7. **Media (M) or Central symmetry system (CSS):** In nymphalids, this band is traditionally treated as two separate bands (M1 and M2) on either side of the discal spot. In papilionids, it appears as a single band, usually positioned near the discal spot, and it is only present on the hindwing.
8. **Umbra (U = r1/r0):** An ambiguous pattern, labeled without description in Schwanwitsch's prototypes. In an earlier work on the nymphalid ground plan, he described it as a shading around the border ocelli (Schwanwitsch, 1924), though it is no longer recognized in the modern nomenclature. Here, we repurpose Umbra in papilionids to represent the orange shading bordering the glauca band. (Schwanwitsch, 1943) referred to this shading as "r0/r1" but did not assign a name for it, believing they may have been Rubrae on the basis of similar coloration.
9. **Externa (E):** The band bordering the distal wing margin. Schwanwitsch recognized two separate bands (E1 and E2), especially in the Parnassiinae. However, we unambiguously identified a single band and are treating it as such for this study.

10. **Glauca (G = E3):** Also defined as Externa 3 (Schwanwitsch, 1956), this prominent dark band runs parallel to the wing margin. Often features an iridescent blue interspace, although it evolved various other states, such as the submarginal chevrons, ocelli, and intervenous stripe.
11. **Submarginal spots (SMS):** The row of spots between G and E. Although such patterns would traditionally be treated as background and not pattern elements (Nijhout, 1991), the SMS are evidently highly prevalent and diverse across Papilionidae. Moreover, they are apparently immune to wing-wide melanization e.g. mimetic *Papilio glaucus* (Perlman and Perlman, 2019)*, Wnt6* RNAi knockdown afffected their size (Iijima et al., 2019) and they are key part of the mimietic forms of *Pachliopta aristolochiae* as *Papilio polytes* and *Papilio alphenor* (Shimajiri and Otaki, 2022)
12. **Marginal spots (MS):** The row of semi-spots bordering the distal wing margin. Like the SMS, these patterns have not traditionally been treated as pattern elements, but they are superficial similarity to the SMS.

### Table S1. List of species and webpage source used in the ancestral reconstruction.

| *Species* | Webpage source |
|---|---|
| *Vanessa cardui (Outgroup)* | https://www.butterfliesofamerica.com/L/imagehtmls/Nymph/Vanessa_cardui_M_USA_PA_Centre_Co_Spring_Creek_ex_larva_05-VII-83-MGCL-2_i.htm |
| | https://www.butterfliesofamerica.com/L/imagehtmls/Nymph/Vanessa_cardui_M_USA_PA_Centre_Co_Spring_Creek_ex_larva_05-VII-83-MGCL-3_i.htm |
| *Baronia brevicornis (non-mimetic)* | https://www.butterfliesofamerica.com/L/imagehtmls/PapPie/Baronia_b_brevicornis_M_MX_GRO_Mexcala_August_1957_MGCL-42143_2_i.htm |
| | https://www.butterfliesofamerica.com/L/imagehtmls/PapPie/Baronia_b_brevicornis_M_MX_GRO_Mexcala_August_1957_MGCL-42143_3_i.htm |
| *Hypermnestra helios* | https://www.europeana.eu/en/item/11622/GLOBISXMFNXGERMANYX6084?q=hypermnestra%20helios&utm_source=old-website&utm_medium=button |
| *Parnassius (Parnassius) apollo* | https://www.europeana.eu/en/item/11622/GLOBISXMFNXGERMANYX5635 |
| *Parnassius (Lingamius) hardwickii* | https://classic.europeana.eu/portal/en/record/11622/GLOBISXMFNXGERMANYX510.html?q=parnassius+acco#dcId=1603707905760&p=1 |
| *Parnassius (Driopa) mnemosyne* | https://www.europeana.eu/en/item/11622/GLOBISXMFNXGERMANYX512 |
| *Parnassius (Kriezbergius) simo* | https://www.europeana.eu/en/item/11622/GLOBISXMFNXGERMANYX5652 |
| *Archon apollinus* | https://www.europeana.eu/en/item/11622/GLOBISXMFNXGERMANYX504 |
| *Archon apollinaris* | https://www.europeana.eu/en/item/11622/GLOBISXMFNXGERMANYX502 |
| *Luehdorfia japonica* | https://www.europeana.eu/en/item/11622/GLOBISXMFNXGERMANYX8515 |
| *Sericinus montela (male)* | https://onlinelibrary.wiley.com/doi/abs/10.1111/azo.12072#support-information-section |
| *Sericinus montela (female)* | https://onlinelibrary.wiley.com/doi/abs/10.1111/azo.12072#support-information-section |
| *Bhutanitis lidderdalii* | http://globis-images.insects-online.de/images/London/London_06/spinosa_Stichel_Bhutanitis_lidderdaleii_BMNH_1A.jpg |
| | http://globis-images.insects-online.de/images/London/London_06/spinosa_Stichel_Bhutanitis_lidderdaleii_BMNH_1B.jpg |
| *Bhutanitis mansfieldi* | http://globis-images.insects-online.de/images/London/London_04/mansfieldi_Riley_Armandia_BMNH_1A.jpg |
| | http://globis-images.insects-online.de/images/London/London_04/mansfieldi_Riley_Armandia_BMNH_1B.jpg |
| *Zerynthia rumina* | https://www.europeana.eu/en/item/11622/GLOBISXMFNXGERMANYX4592 |
| *Allancastria cerisyi* | http://insecta.pro/taxonomy/46120 |
| *Lamproptera meges* | http://insecta.pro/gallery/35545 |
| *Iphiclides podalirius* | https:://www.europeana.eu/en/item/11622/GLOBISXMFNXGERMANYX4038 |
| *Protographium marcellus* | https://www.butterfliesofamerica.com/L/imagehtmls/PapPie/31_Eurytides_marcellus_F__USA__TEXAS__Tyler_Co__Kirby_S_F__19-IV-94___marctelefd_i.htm |
| | https://www.butterfliesofamerica.com/L/imagehtmls/PapPie/32_Eurytides_marcellus_F__USA__TEXAS__Tyler_Co__Kirby_S_F__19-IV-94___martelefv_i.htm |
| *Mimoides thymbraeus* | https:://www.butterfliesofamerica.com/L/imagehtmls/PapPie/Mimoides_t_thymbraeus_M_GUATEMALA_June_1916_2_i.htm |
| | https:://www.butterfliesofamerica.com/L/imagehtmls/PapPie/Mimoides_t_thymbraeus_M_GUATEMALA_June_1916_3_i.htm |
| *Eurytides dolicaon* | https:://www.butterfliesofamerica.com/L/imagehtmls/PapPie/Eurytides_dolicaon_septentrionalis_M_PANAMA_CZ_Madden_Dam_08-V-66_2_i.htm |
| | https:://www.butterfliesofamerica.com/L/imagehtmls/PapPie/Eurytides_dolicaon_septentrionalis_M_PANAMA_CZ_Madden_Dam_08-V-66_3_i.htm |
| *Protesilaus protesilaus* | https:://www.butterfliesofamerica.com/L/imagehtmls/PapPie/Eurytides_protesilaus_dariensis_M_PANAMA_2_i.htm |
| | https:://www.butterfliesofamerica.com/L/imagehtmls/PapPie/Eurytides_protesilaus_dariensis_M_PANAMA_3_i.htm |
| *Graphium (Pathysa) delesserti* | http://insecta.pro/gallery/38349 |
| | http://insecta.pro/gallery/38350 |
| *Graphium (Arisbe) leonidas* | https:://www.europeana.eu/en/item/11623/_ENBIBUTTERFLIES_MRAC_BELGIUM_740030330_mr |
| | https:://www.europeana.eu/en/item/11623/_ENBIBUTTERFLIES_MRAC_BELGIUM_740030330_mv |
| *Graphium (Pazala) incertus* | https:://www.europeana.eu/en/item/11622/GLOBISXMFNXGERMANYX5508 |
| *Graphium (Idaides) sarpedon* | http://insecta.pro/gallery/38596 |
| | http://insecta.pro/gallery/38597 |
| *Teinopalpus imperialis* | https:://www.aureus-butterflies.de/epages/17926950.sf/en_GB/?ObjectID=68626804 |
| *Teinopalpus aureus* | http://insecta.pro/gallery/26881 |
| | http://insecta.pro/gallery/26882 |
| *Meandrusa payeni* | http://insecta.pro/gallery/32425 |
| | http://insecta.pro/gallery/32426 |
| *Meandrusa sciron* | http://insecta.pro/gallery/32423 |
| | http://insecta.pro/gallery/32424 |
| *Papilio (Heraclides) cresphontes* | https:://www.butterfliesofamerica.com/L/imagehtmls/PapPie/Papilio_cresphontes_f1d2_USA_FLORIDA_Broward_Co._Ft._Lauderdale_emg._18-VIII-1978-MGCL_i.htm |
| | https:://www.butterfliesofamerica.com/L/imagehtmls/PapPie/Papilio_cresphontes_f1v2_USA_FLORIDA_Broward_Co._Ft._Lauderdale_emg._18-VIII-1978-MGCL_i.htm |
| *Papilio (Heraclides) machaonides* | https:://www.butterfliesofamerica.com/L/imagehtmls/PapPie/Papilio_machaonides_M_DR_BARAHONA_PROV_Rio_Baoruco_Canyon_10_km_S_of_Barahona_50-350_17-IX-1995-MGC_L-2_i.htm |
| | https:://www.butterfliesofamerica.com/L/imagehtmls/PapPie/Papilio_machaonides_M_DR_BARAHONA_PROV_Rio_Baoruco_Canyon_10_km_S_of_Barahona_50-350_17-IX-1995-MGC_L-3_i.htm |
| *Papilio (Heraclides) anchisiades* | https:://www.butterfliesofamerica.com/L/imagehtmls/PapPie/04-SRNP-41288-DHJ97404_i.htm |
| | https:://www.butterfliesofamerica.com/L/imagehtmls/PapPie/04-SRNP-41288-DHJ97405_i.htm |
| *Papilio (Heraclides) astyalus (non-mimetic)* | https:://www.butterfliesofamerica.com/L/imagehtmls/PapPie/Papilio_astyalus_hippomedon_M_VENEZUELA_25_km_S_Bolivar_El_Dorado_2_i.htm |
| | https:://www.butterfliesofamerica.com/L/imagehtmls/PapPie/Papilio_astyalus_hippomedon_M_VENEZUELA_25_km_S_Bolivar_El_Dorado_3_i.htm |
| *Papilio (Heraclides) astyalus (mimetic)* | https:://www.butterfliesofamerica.com/L/imagehtmls/PapPie/Papilio_astyalus_hippomedon_F_TRINIDAD_1926_2_i.htm |
| | https:://www.butterfliesofamerica.com/L/imagehtmls/PapPie/Papilio_astyalus_hippomedon_F_TRINIDAD_1926_3_i.htm |
| *Papilio (Alexanoria) alexanor* | https:://www.europeana.eu/en/item/11622/GLOBISXMFNXGERMANYX501 |
| *Papilio (Chilasa) clytia* | https:://www.aureus-butterflies.de/epages/17926950.sf/en_GB/?ObjectPath=/Shops/17926950/Products/%22Papilio%20clytia%20dissimilis%20M%C3%A4nnchen%22 |
| *Papilio (Chilasa) moerneri* | https:://www.aureus-butterflies.de/epages/17926950.sf/en_GB/?ObjectPath=/Shops/17926950/Products/%22Chilasa%20moerneri%20moerneri%20M%22 |
| *Papilio (Pterourus) esperanza* | https:://www.butterfliesofamerica.com/L/imagehtmls/PapPie/Papilio_esperanza_HOLOTYPE_M_[IBUNAM]_2_i.htm |

### Continuation Table S1.

| | |
|---|---|
| | https:://www.butterfliesofamerica.com/L/imagehtmls/PapPie/Papilio_esperanza_HOLOTYPE_M_[IBUNAM]_3_i.htm |
| *Papilio (Pterourus) troilus* | https:://www.butterfliesofamerica.com/L/imagehtmls/PapPie/Papilio_troilus_NEOTYPE_2_i.htm<br>https:://www.butterfliesofamerica.com/L/imagehtmls/PapPie/Papilio_troilus_NEOTYPE_3_i.htm |
| *Papilio (Pterourus) palamedes* | https:://www.butterfliesofamerica.com/L/imagehtmls/PapPie/61_Papilio_p_palamedes_Springfeild_Bay_Co_FL_USA_11-VI-78_1_i.htm<br>https:://www.butterfliesofamerica.com/L/imagehtmls/PapPie/62_Papilio_p_palamedes_Springfeild_Bay_Co_FL_USA_11-VI-78_2_i.htm |
| *Papilio (Pterourus) pilumnus* | https:://www.butterfliesofamerica.com/L/imagehtmls/PapPie/Papilio_pilumnus_M_Mixcoate_MX_10-IX-04_1_i.htm<br>https:://www.butterfliesofamerica.com/L/imagehtmls/PapPie/Papilio_pilumnus_M_Mixcoate_MX_10-IX-04_2_i.htm |
| *Papilio (Agehana [Pterourus]) maraho* | https:://www.aureus-butterflies.de/epages/17926950.sf/en_GB/?ObjectPath=/Shops/17926950/Products/%22Agehana%20maraho%20Weibchen%22 |
| *Papilio (Pterourus) glaucus (non-mimetic)* | https:://www.butterfliesofamerica.com/L/imagehtmls/PapPie/Papilio_g_glaucus_NEOTYPE_M_2_i.htm<br>https:://www.butterfliesofamerica.com/L/imagehtmls/PapPie/Papilio_g_glaucus_NEOTYPE_M_3_i.htm |
| *Papilio (Pterourus) glaucus (mimetic)* | https:://www.butterfliesofamerica.com/L/imagehtmls/PapPie/Papilio_g_glaucus_NEOTYPE_F_5_i.htm<br>https:://www.butterfliesofamerica.com/L/imagehtmls/PapPie/Papilio_g_glaucus_NEOTYPE_F_6_i.htm |
| *Papilio (Pterourus) zagreus* | https:://www.butterfliesofamerica.com/L/ih/DSCN8180_i.htm<br>https:://www.butterfliesofamerica.com/L/ih/DSCN8181_i.htm |
| *Papilio (Pterourus) garamas (non-mimetic)* | https:://www.butterfliesofamerica.com/L/imagehtmls/PapPie/Papilio_garamas_NW_Mexican_segregate_M_MX_NAJ_La_Yerba_09-IX-80_MZFC_3_i.htm<br>https:://www.butterfliesofamerica.com/L/imagehtmls/PapPie/Papilio_garamas_NW_Mexican_segregate_M_MX_NAJ_La_Yerba_09-IX-80_MZFC_2_i.htm |
| *Papilio (Eleppone) anactus* | https:://www.aureus-butterflies.de/epages/17926950.sf/en_GB/?ObjectPath=/Shops/17926950/Products/%22Papilio%20anactus%20ssp.%20%3F%20M%C3%A4nnchen%22 |
| *Papilio (Druryia [Princeps]) antimachus* | https:://www.aureus-butterflies.de/epages/17926950.sf/en_GB/?ObjectPath=/Shops/17926950/Products/%22Papilio%20antimachus%20antimachus%20M%C3%A4nnchen%22 |
| *Papilio (Sinoprinceps) xuthus* | http://insecta.pro/gallery/38580<br>http://insecta.pro/gallery/38579 |
| *Papilio (Papilio) zelicaon* | https:://www.butterfliesofamerica.com/L/imagehtmls/PapPie/Papilio_zelicaon_M_Edison_Kern_Co_CA_USA_25-VI-73_C2_2_i.htm<br>https:://www.butterfliesofamerica.com/L/imagehtmls/PapPie/Papilio_zelicaon_M_Edison_Kern_Co_CA_USA_25-VI-73_C2_3_i.htm |
| *Papilio (Papilio) polyxenes* | https:://www.butterfliesofamerica.com/L/imagehtmls/PapPie/Papilio_polyxenes_asterius_M_Tamazunchale_SLP_MX_4-VII-87_152_i.htm<br>https:://www.butterfliesofamerica.com/L/imagehtmls/PapPie/Papilio_polyxenes_asterius_M_Tamazunchale_SLP_MX_4-VII-87_154_i.htm |
| *Papilio (Druryia [Princeps]) zalmoxis* | https:://www.aureus-butterflies.de/epages/17926950.sf/en_GB/?ObjectPath=/Shops/17926950/Products/%22Papilio%20zalmoxis%20M%C3%A4nnchen%22 |
| *Papilio (Druryia [Princeps]) nireus* | https:://www.aureus-butterflies.de/epages/17926950.sf/en_GB/?ObjectPath=/Shops/17926950/Products/%22Papilio%20nireus%20nireus%20M%C3%A4nnchen%22 |
| *Papilio (Achillides) ulysses* | https:://www.aureus-butterflies.de/epages/17926950.sf/en_GB/?ObjectPath=/Shops/17926950/Products/%22Papilio%20ulysses%20ulysses%20M%C3%A4nnchen%22 |
| *Papilio (Achillides) maackii* | https:://www.aureus-butterflies.de/epages/17926950.sf/en_GB/?ObjectPath=/Shops/17926950/Products/%22Papilio%20maackii%20tutanus%20Weibchen%22 |
| *Papilio (Princeps) demoleus* | https:://www.aureus-butterflies.de/epages/17926950.sf/en_GB/?ObjectPath=/Shops/17926950/Products/%22Papilio%20demoleus%20ssp.%20%3F%20M%C3%A4nnchen%22 |
| *Papilio (Menelaides) polytes (non-mimetic)* | https:://www.aureus-butterflies.de/epages/17926950.sf/en_GB/?ObjectPath=/Shops/17926950/Products/%22Papilio%20polytes%20alphenor%20M%C3%A4nnchen%22 |
| *Papilio (Menelaides) memnon* | https:://www.aureus-butterflies.de/epages/17926950.sf/en_GB/?ObjectPath=/Shops/17926950/Products/%22Papilio%20memnon%20memnon%20M%C3%A4nnchen%22 |
| *Battus philenor* | https:://www.butterfliesofamerica.com/L/imagehtmls/PapPie/6_Battus_philenor_philenor_M_Florida_Wash_Madera_Cyn_Pima_Co_AZ_USA_4-IX-78_C2_0049_i.htm<br>https:://www.butterfliesofamerica.com/L/imagehtmls/PapPie/6_Battus_philenor_philenor_M_Florida_Wash_Madera_Cyn_Pima_Co_AZ_USA_4-IX-78_C2_0049_1_i.htm |
| *Battus polydamas* | https:://www.butterfliesofamerica.com/L/imagehtmls/PapPie/Battus_polydamas_polydamas_Cd_Victoria_TAMP_MX_23-IV-87_013_DR_Coll_i.htm<br>https:://www.butterfliesofamerica.com/L/imagehtmls/PapPie/Battus_polydamas_polydamas_Cd_Victoria_TAMP_MX_23-IV-87_015_DR_Coll_i.htm |
| *Battus crassus* | https:://www.butterfliesofamerica.com/images/Papilionidae/Papilioninae/Battus_crassus_lepidus/Battus_crassus_lepidus_M_Colombia_0029.jpg<br>https:://www.butterfliesofamerica.com/images/Papilionidae/Papilioninae/Battus_crassus_lepidus/Battus_crassus_lepidus_M_Colombia_0030.jpg |
| *Pharmacophagus antenor* | http://insecta.pro/gallery/31908<br>http://insecta.pro/gallery/31906 |
| *Cressida cressida* | https:://www.aureus-butterflies.de/epages/17926950.sf/en_GB/?ObjectPath=/Shops/17926950/Products/%22Cressida%20cressida%20cassandra%20M%C3%A4nnchen%22 |
| *Byasa polyeuctes* | https:://www.aureus-butterflies.de/epages/17926950.sf/en_GB/?ObjectPath=/Shops/17926950/Products/%22Byasa%20polyeuctes%20M%C3%A4nnchen%22 |
| *Atrophaneura semperi* | https:://www.aureus-butterflies.de/epages/17926950.sf/en_GB/?ObjectPath=/Shops/17926950/Products/%22Atrophaneura%20semperi%20Weibchen%22 |
| *Atrophaneura horishanus* | https:://www.aureus-butterflies.de/epages/17926950.sf/en_GB/?ObjectID=70240043 |
| *Losaria neptunus* | https:://www.aureus-butterflies.de/epages/17926950.sf/en_GB/?ObjectPath=/Shops/17926950/Products/%22Losaria%20neptunus%20decasini%20Weibchen%22 |
| *Pachliopta aristolochiae* | https:://www.aureus-butterflies.de/epages/17926950.sf/en_GB/?ObjectPath=/Shops/17926950/Products/%22Pachliopta%20aristolochiae%20aristolochiae%20M%C3%A4nnchen%22 |
| *Euryades duponchelii* | https:://www.aureus-butterflies.de/epages/17926950.sf/en_GB/?ObjectPath=/Shops/17926950/Products/%22Euryades%20duponchelii%20M%C3%A4nnchen%22 |
| *Parides montezuma* | https:://www.butterfliesofamerica.com/L/imagehtmls/PapPie/Parides_montezuma_Guadelajara_Jalisco_MX_11-VIII-96_085_i.htm<br>https:://www.butterfliesofamerica.com/L/imagehtmls/PapPie/Parides_montezuma_Guadelajara_Jalisco_MX_11-VIII-96_086_i.htm |
| *Parides eurimedes* | https:://www.butterfliesofamerica.com/L/imagehtmls/PapPie/Parides_eurimedes_mylotes_M_CR_HEREDIA_Rd_to_Tirimbina_5-7_KM_E_of_Ruta_9_21-III-892_i.htm<br>https:://www.butterfliesofamerica.com/L/imagehtmls/PapPie/Parides_eurimedes_mylotes_M_CR_HEREDIA_Rd_to_Tirimbina_5-7_KM_E_of_Ruta_9_21-III-893_i.htm |
| *Parides photinus* | https:://www.butterfliesofamerica.com/L/imagehtmls/PapPie/Parides_photinus_F_MX_CHIS_Santa_Rosa_Comitan_III-58_2_i.htm<br>https:://www.butterfliesofamerica.com/L/imagehtmls/PapPie/Parides_photinus_F_MX_CHIS_Santa_Rosa_Comitan_III-58_3_i.htm |
| *Parides erithalion* | https:://www.butterfliesofamerica.com/L/imagehtmls/PapPie/Parides_erithalion_trichopus_M_MEXICO_GUERRERO_Acahuizotla_September_1966-MGCL-2_i.htm<br>https:://www.butterfliesofamerica.com/L/imagehtmls/PapPie/Parides_erithalion_trichopus_M_MEXICO_GUERRERO_Acahuizotla_September_1966-MGCL-3_i.htm |
| *Trogonoptera brookiana (male)* | https:://www.aureus-butterflies.de/epages/17926950.sf/en_GB/?ObjectPath=/Shops/17926950/Products/%22Trogonoptera%20brookiana%20albescens%20M%22 |
| *Trogonoptera brookiana (female)* | https:://www.aureus-butterflies.de/epages/17926950.sf/en_GB/?ObjectPath=/Shops/17926950/Products/%22Trogonoptera%20brookiana%20brookiana%20Weibchen%22 |
| *Troides helena (male)* | https:://www.aureus-butterflies.de/epages/17926950.sf/en_GB/?ObjectPath=/Shops/17926950/Products/%22Troides%20helena%20hephaestus%20M%C3%A4nnchen%22 |
| *Troides helena (female)* | https:://www.aureus-butterflies.de/epages/17926950.sf/en_GB/?ObjectPath=/Shops/17926950/Products/%22Troides%20helena%20helena%20f.%20Weibchen%22 |
| *Ornithoptera priamus (male)* | https:://www.aureus-butterflies.de/epages/17926950.sf/en_GB/?ObjectPath=/Shops/17926950/Products/%22Ornithoptera%20priamus%20priamus%20M%C3%A4nnchen%22 |

**Table S2. Species population source and rearing information. All the adults were hand feed with 10% sugar water solution.**

| Species | Population origin | Collected | Maiting | Caterpillar host plant |
|---|---|---|---|---|
| *Papilio zelicaon* | Albany, CA | Yiwen Zhu. Collected eggs and larvae on fennel | Hand paired ~10-20 females per generation | Fennel (*Foeniculum vulgare*)<br>Parsley (*Petroselinum crispum*) |
| *Papilio polyxenes* | Ithaca, NY | Alan Liang, Brian Liang. Collected adult female and larvae on parsley | Hand paired ~10-20 females per generation | Fennel (*Foeniculum vulgare*)<br>Parsley (*Petroselinum crispum*) |
| | South Carolina | Purchased twelve pupae from Kevin Koffel | | |
| *Papilio glaucus* | Iowa | purchased ~180 eggs from David Thompson | Hand paired 3 females for second generation | Tuliptree (*Liriodendron tulipifera)*<br>Chokecherry (*Prunus virginiana*)<br>Ash (*Fraxinus sp.*) |
| *Papilio troilus* | Ithaca, NY | Alan Liang, Brian Liang (collected larva on spicebush) | Hand paired 2 females | Spicebush (*Lindera benzoin*) |
| | Allegan, MI | Purchased ten pupae from Kevin Collison | | |
| *Papilio cresphontes* | Ithaca, NY | Alan Liang, Brian Liang. Collected adult females and eggs on rue and amur cork tree | No mating | Rue (*Ruta graveolens*) |
| *Protographium marcellus* | Columbia District | Arnaud Martin, Anyi Mazo-Vargas. Collected eggs, larvae on pawpaw trees | No mating | Pawpaw tree (*Asimina triloba*) |

**Table S3. Heparin and Dextran sulfate drug injections information.**

| Species | Age of pupae: Hours after pupation | Treatment | Amount injected (ug) | Optimal* amount (ug) | Pupae injected | Adults with phenotype |
|---|---|---|---|---|---|---|
| *P. zelicaon* | 4-20 | Heparin | 60-200 | 80-120 | ~20 | 10 |
| *P. polyxenes* | 4-20 | Heparin | 60-200 | 80-120 | 30-40 | 16 |
| *P. glaucus* | 4-24 | Heparin | 60-320 | >120 | ~30 | 6 |
| *P. troilus* | 4-10 | Heparin | 60-160 | 100-120 | ~15 | 2 |
| *P. cresphontes* | 4-10 | Heparin | 60-160 | 120 | ~20 | 1 |
| *P. zelicaon* | 2-10 | Dextran sulfate | 20-60 | 20-60 | ~20 | 13 |
| *P. polyxenes* | 4-20 | Dextran sulfate | 20-100 | ? | 20-30 | 0 |
| *P. glaucus* | 4-20 | Dextran sulfate | 20-200 | ? | ~30 | 0 |
| *P. troilus* | 4-10 | Dextran sulfate | 20-100 | ? | ~10 | 0 |
| *P. cresphontes* | 4-10 | Dextran sulfate | 20-100 | ? | ~20 | 0 |

**Table S4. List of oligos used for DNA amplification and sgRNA used in the CRISPR-Cas9 knockout experiments.**

| **Oligo** | **sequence** | **Function** | **gene** |
|---|---|---|---|
| Papilio_WntApr1_FWD | GATGGTGGCCCATGGTAGG | PCR oligo | *WntA* |
| Papilio_WntApr1_REV | CCACCAATTACCGGTGACAGC | PCR oligo | *WntA* |
| Papilio_WntA_Fwd_ISH | GCACTGGCAATGGGGAGG | PCR oligo | *WntA* |
| Papilio_WntA_REV_ISH | GCAATGTACTCGACAGCACC | PCR oligo | *WntA* |
| Papilio_wnt6_Ex3_FWD | GCTGGGGTGACATACGCCAT | PCR oligo | *Wnt6* |
| Papilio_wnt6_Ex6_REV | GGTCACAACTATCCGTTCCGGC | PCR oligo | *Wnt6* |
| NP_PmWtnA_FWD | GTGCAAATCGCGGTGCAAATC | PCR oligo | *WntA* |
| NP_PmWtnA_REV | GCACGTCCAGGAACTGTAAACC | PCR oligo | *WntA* |
| NP_PmWtn6_FWD | CGAGGCGTACCAATGACCAC | PCR oligo | *Wnt6* |
| NP_PmWtn6_REV | ATCGAACCCAGGAAGAAGCC | PCR oligo | *Wnt6* |
| **gRNA** | | | |
| PzPp_WntApr_sgRNA1 | AAGGGGAAGGGTCGTACTGG | sgRNA | *WntA* |
| PzPp_WntApr_sgRNA2 | ATTCGAAAGAGACTTTGGAA | sgRNA | *WntA* |
| PzPp_WntApr_sgRNA3 | AAGTGATTTAAATACACAAT | sgRNA | *WntA* |
| PzPp_WntAE1_sgRNA4 | AACTTTCCAAAATAGCTTCA | sgRNA | *WntA* |
| Pz_Wnt6_sgRNA1 | CATCACACGAGCCTGCACTG | sgRNA | *Wnt6* |
| Pz_Wnt6_sgRNA2 | CATCACACGAGCCTGCACTG | sgRNA | *Wnt6* |

**Table S5. *WntA* and *Wnt6* CRISPR-Cas9 injections information.**

| Species | Gene | Time of egg collection (hours) | eggs injected | Number of L1 larvae | Hatching Ratio | Number of pupae | Number of adults with phenotype | Phenotype ratio from hatched eggs |
|---|---|---|---|---|---|---|---|---|
| *P. zelicaon* | *WntA* | 1-4 | ~400 | 60 | 15% | 12 | 5 | 8% |
| *P. polyxenes* | *WntA* | 1-4 | ~520 | 81 | 16% | 19 | 5 | 14% |
| *P. zelicaon* | *Wnt6* | 1-4 | ~190 | 35 | 18% | 13 | 7 | 20% |
| *P. polyxenes* | *Wnt6* | 1-4 | ~350 | 25 | 7% | 9 | 6 | 24% |

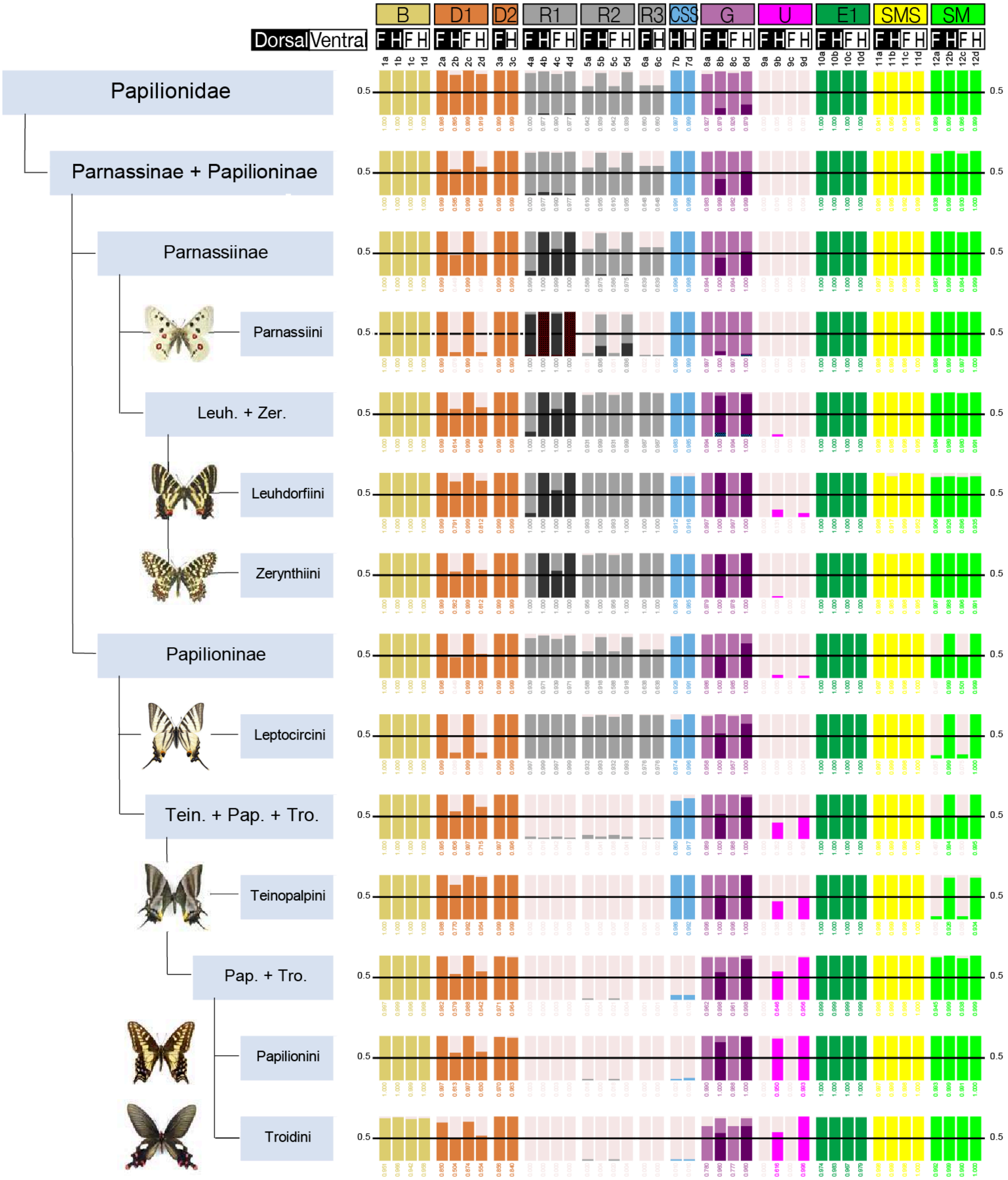


**Figure S1.** Relative ancestral state likelihoods for major papilionid clades represented by stacked bar charts.

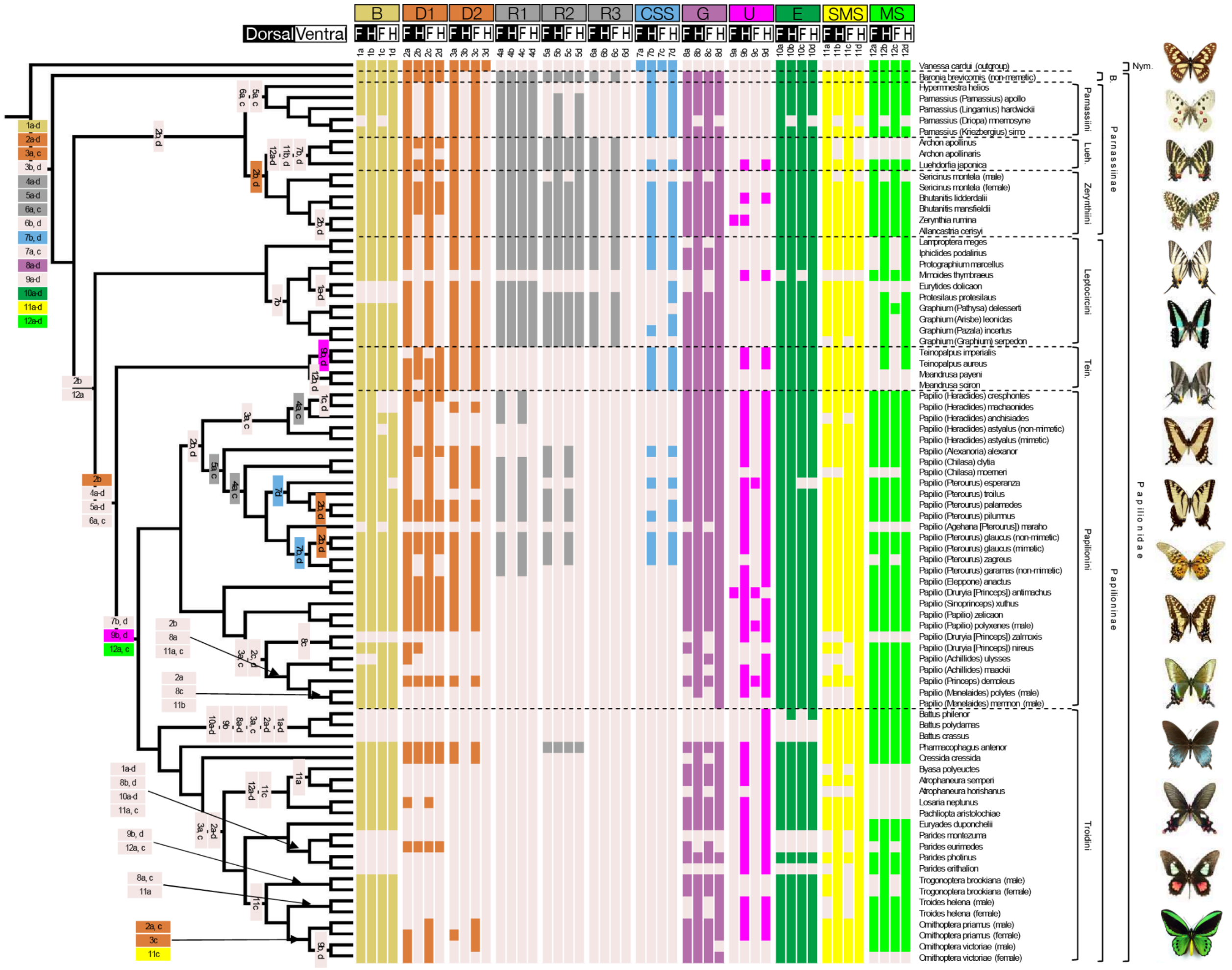


**Figure S2.** Distribution of pattern element character states (presence/absence) across papilionid species. The colored boxes at the Papilionidae node represent ancestral character states of the family based on ML methods. Subsequent state changes are shown as colored boxes at lower nodes.

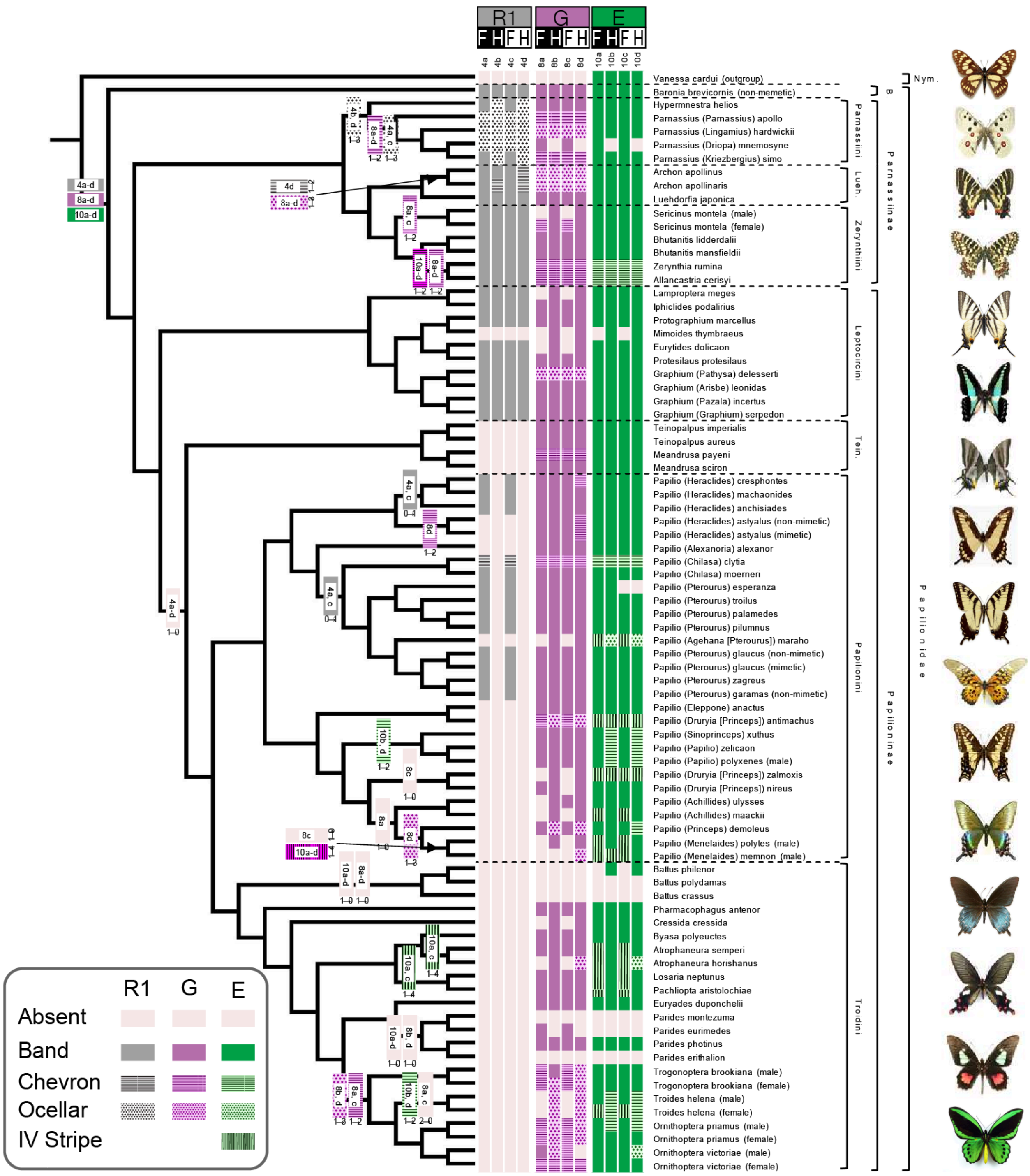


**Figure S3.** Distribution of shape states observed in pattern elements R1, G, and E across papilionid species. The colored boxes at the Papilionidae node represent ancestral character states of the family based on ML methods. Subsequent state changes are shown as colored boxes at lower nodes.

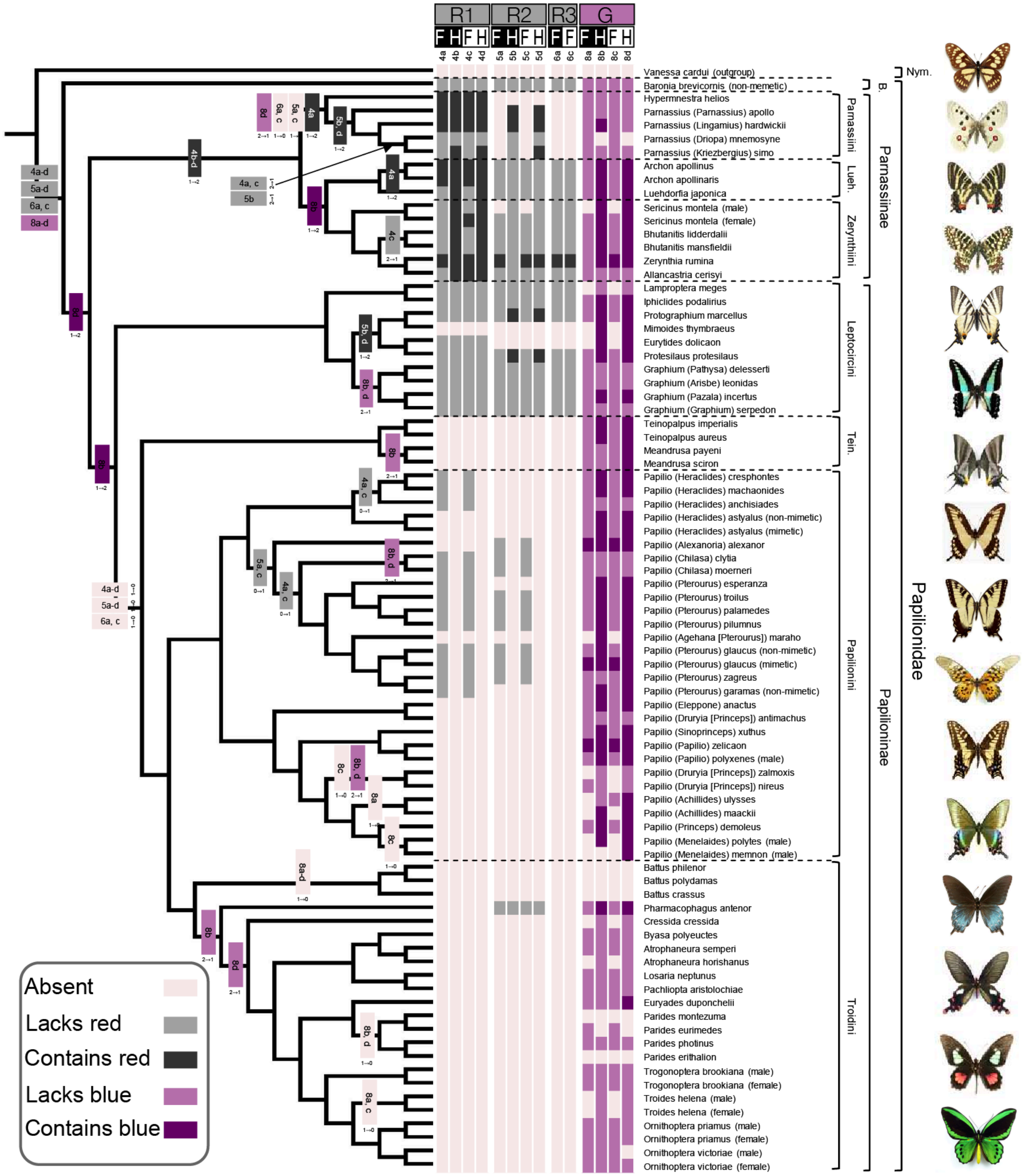


**Figure S4.** Distribution of color states observed in pattern elements R1, R2, R3, and G across papilionid species. The colored boxes at the Papilionidae node represent ancestral character states of the family based on ML methods. Subsequent state changes are shown as colored boxes at lower nodes.

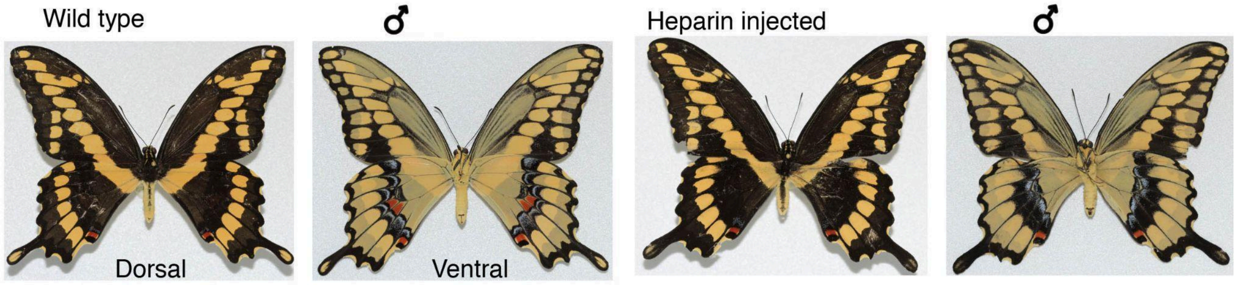


**Figure S5.** Effects of heparin on *Papilio cresphontes*.

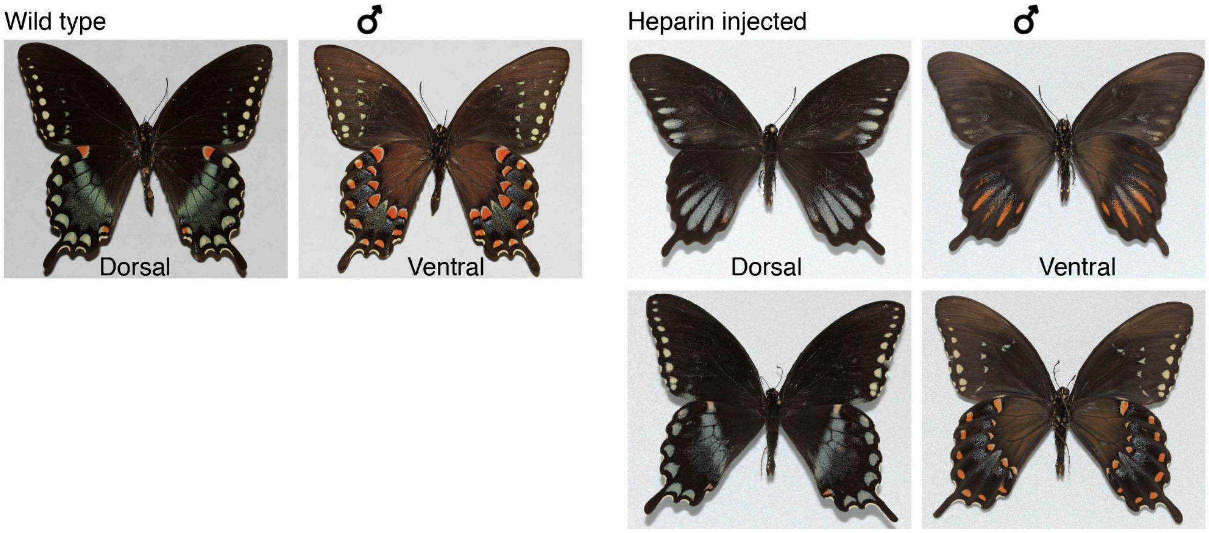


**Figure S6.** Effects of heparin on *Papilio trolius*.

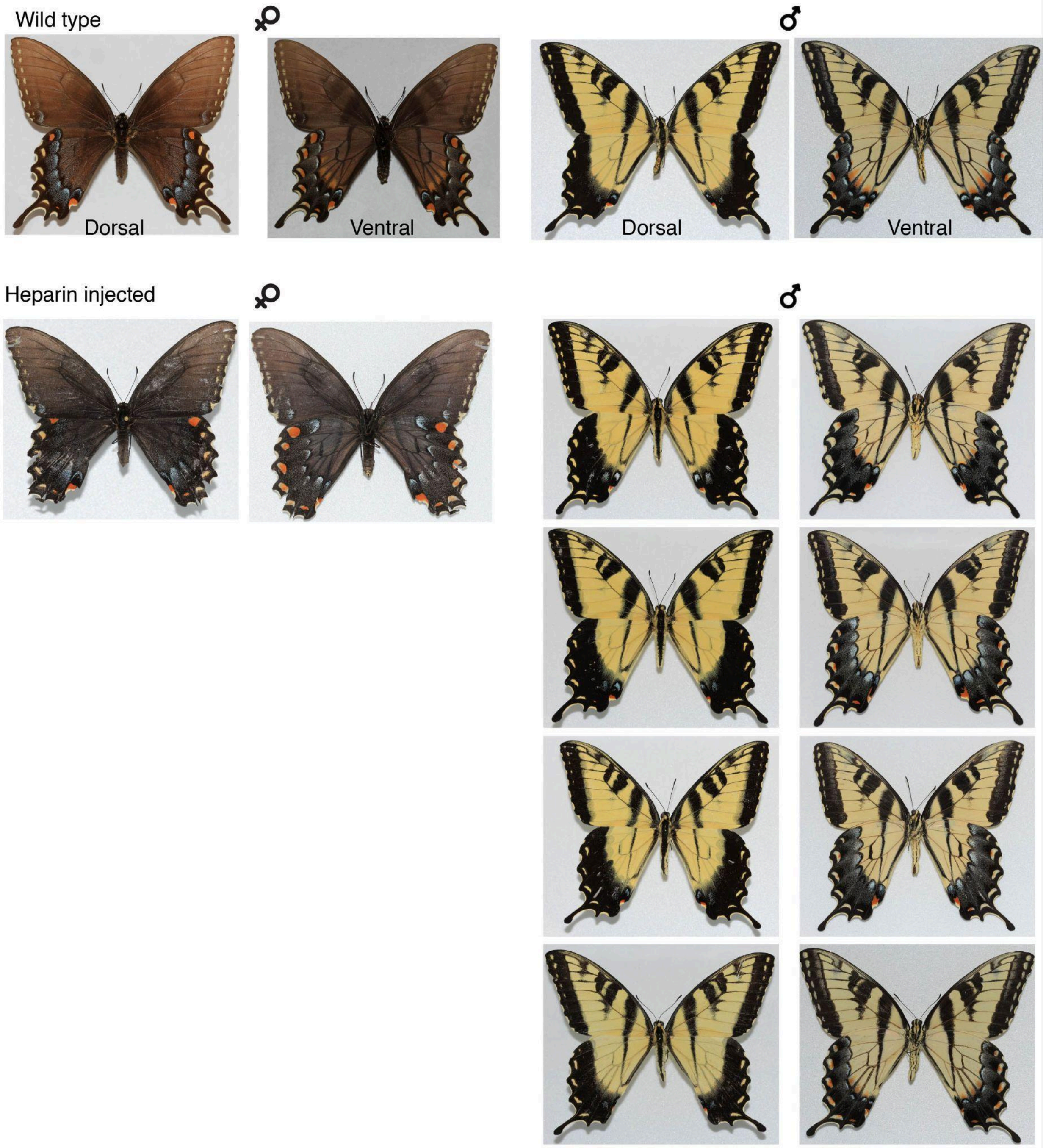


**Figure S7.** Effects of heparin on *Papilio glaucus*.

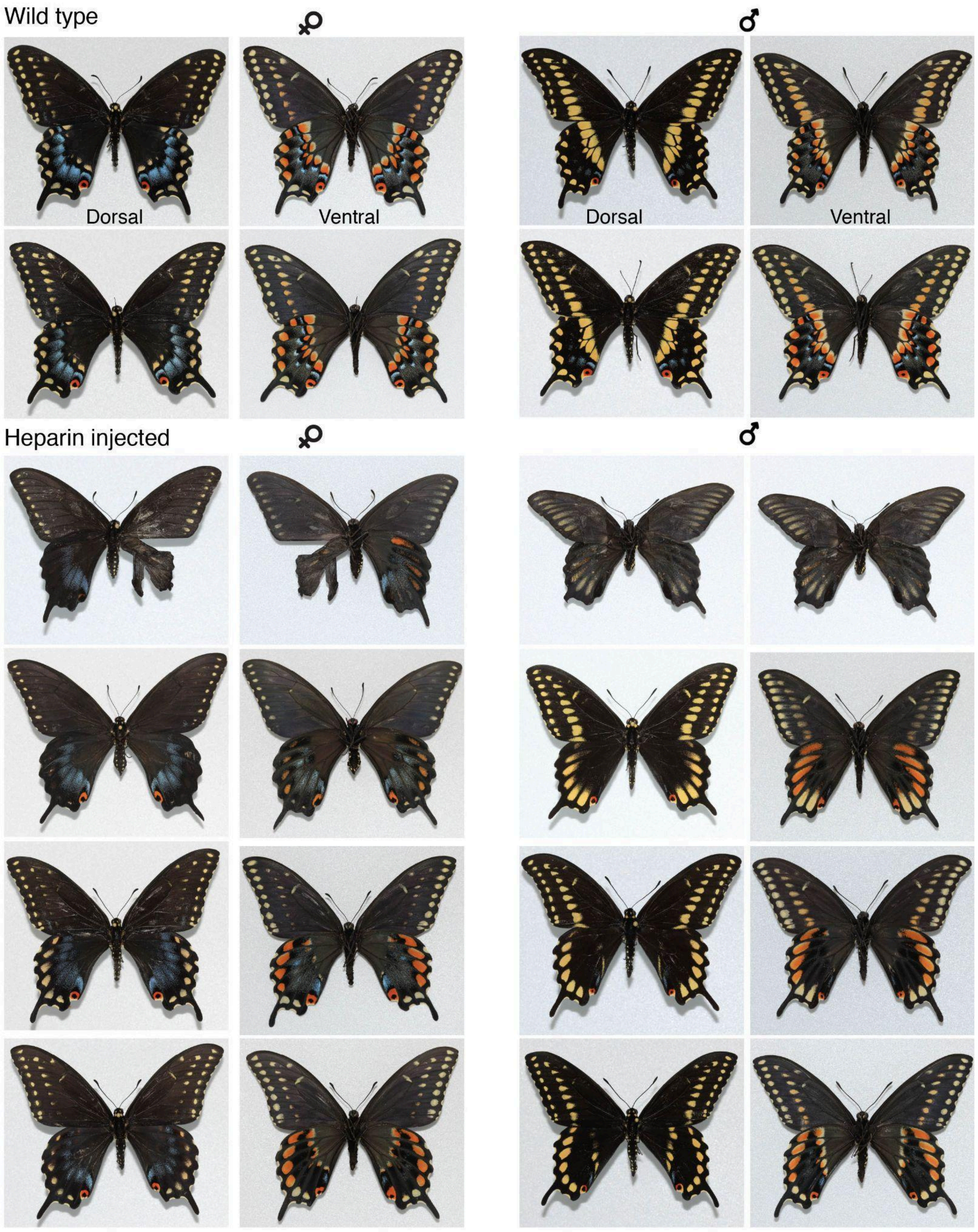


**Figure S8.** Effects of heparin on *Papilio polyxenes*.

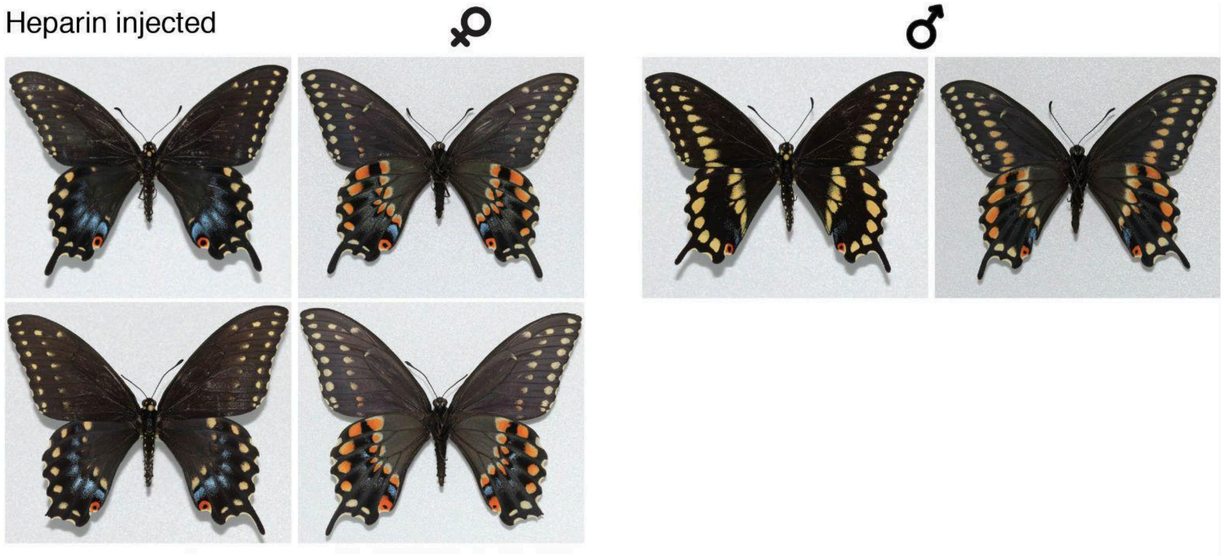


**Continuation Figure S8.** Effects of heparin on *Papilio polyxenes*.

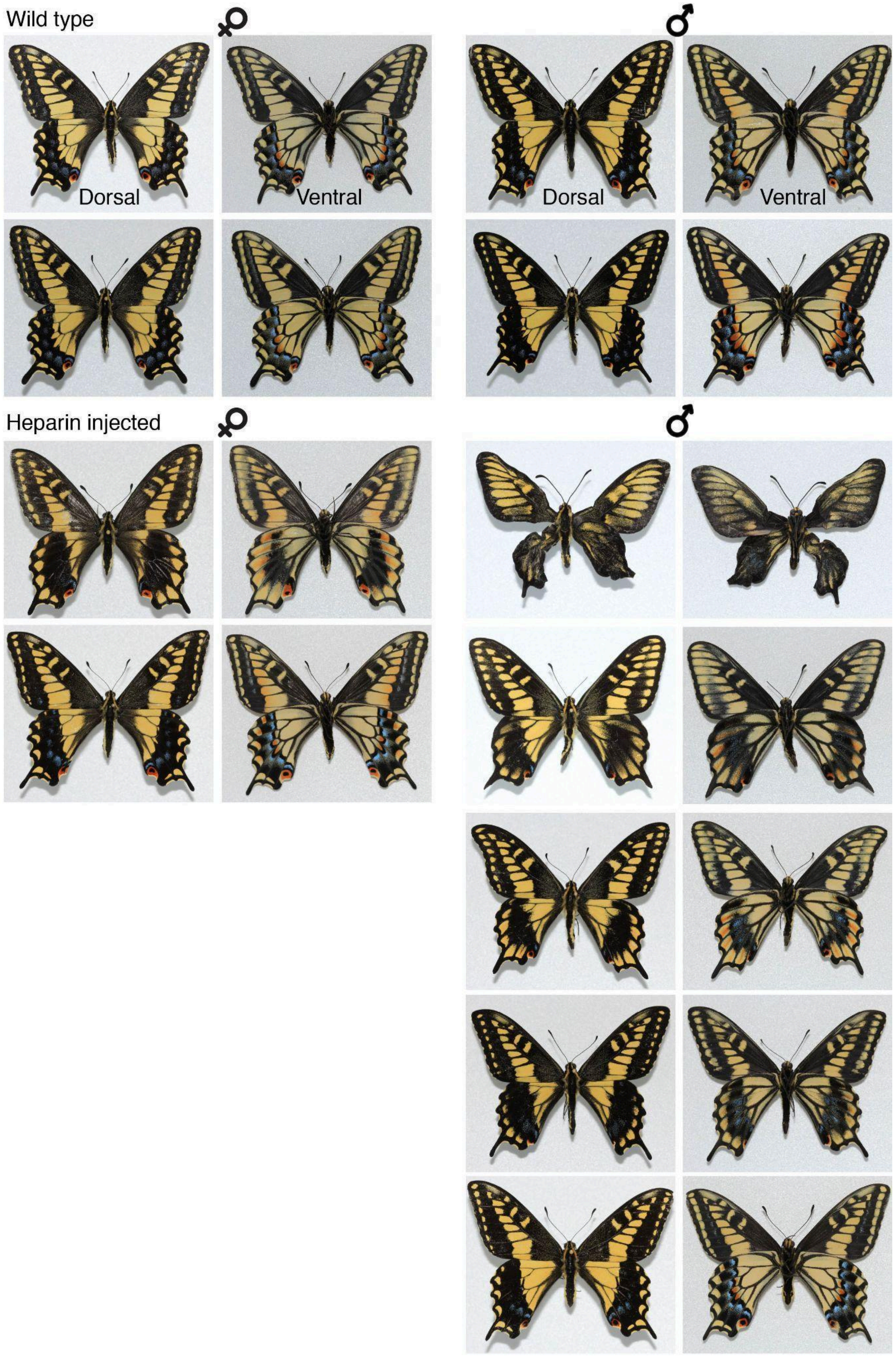

Wild type
♀
♂
Dorsal
Ventral
Dorsal
Ventral
Heparin injected
♀
♂

**Figure S9.** Effects of heparin on *Papilio zelicaon*.

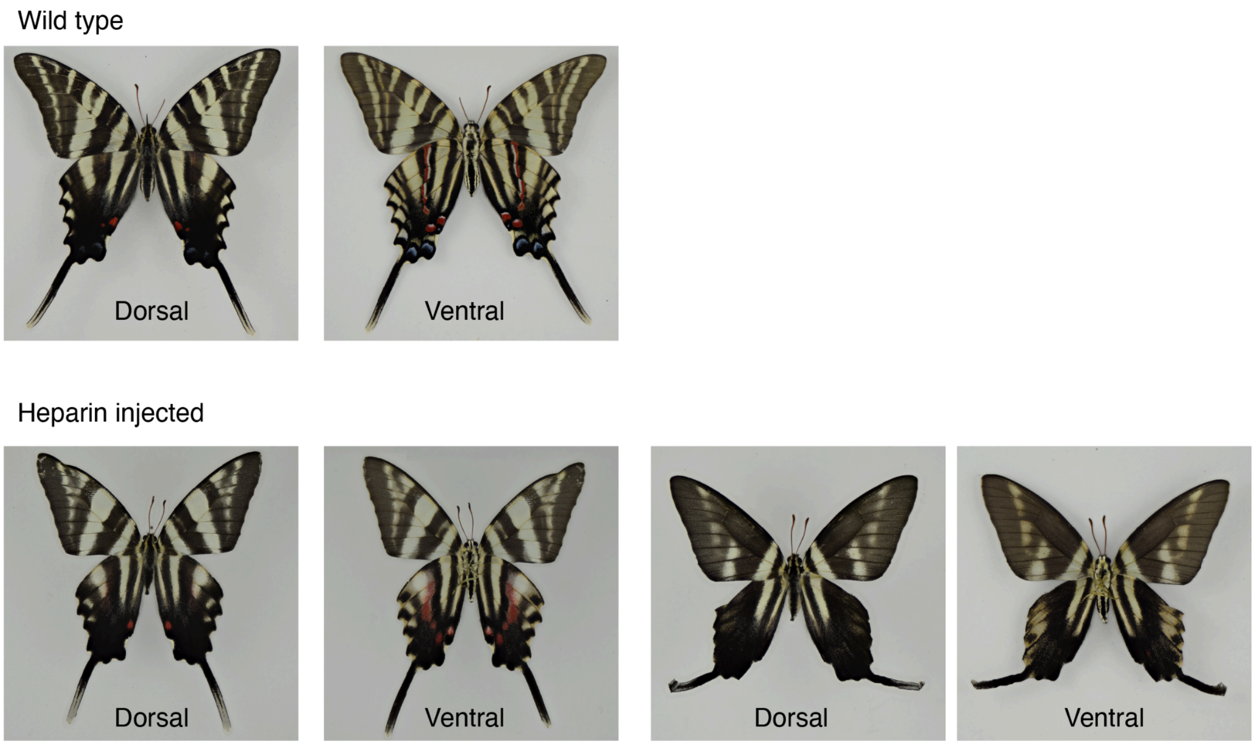


**Figure S10.** Effects of heparin on *Protographium marcellus*.

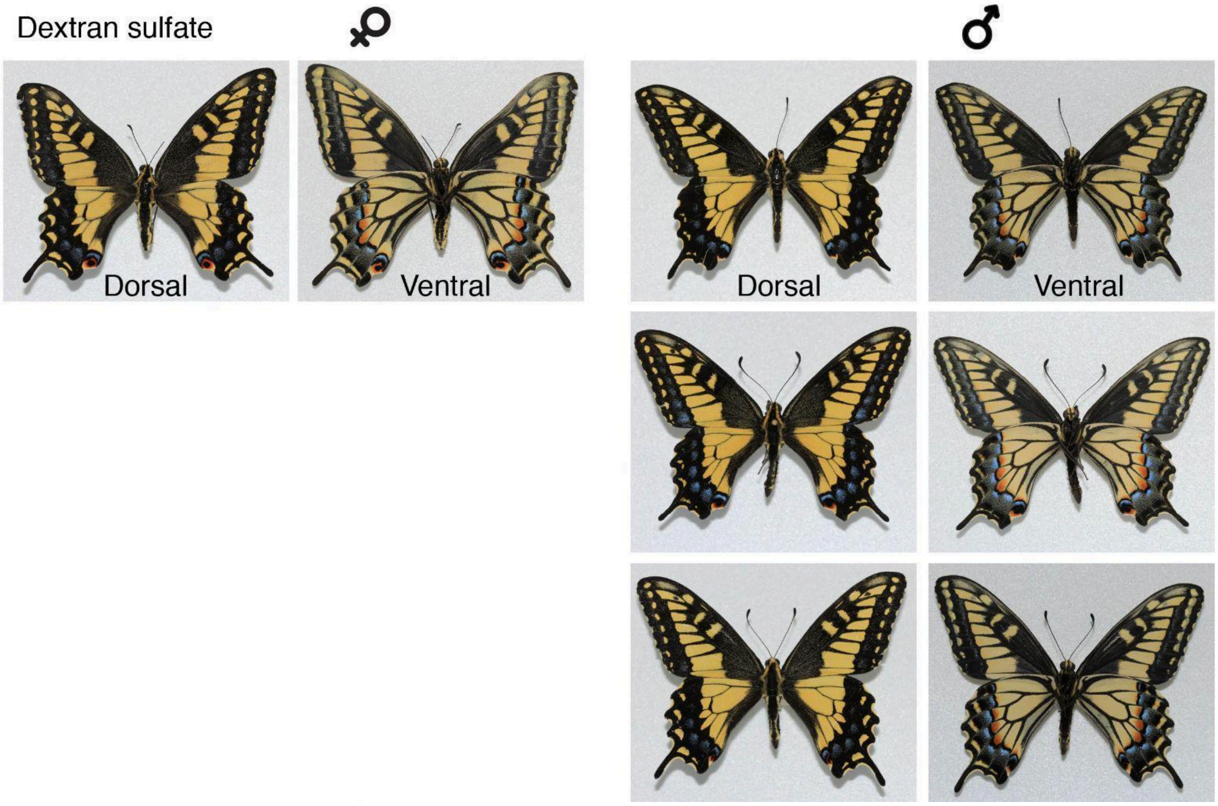


**Figure S11.** Effects of Dextran sulfate on *Papilio zelicaon*.

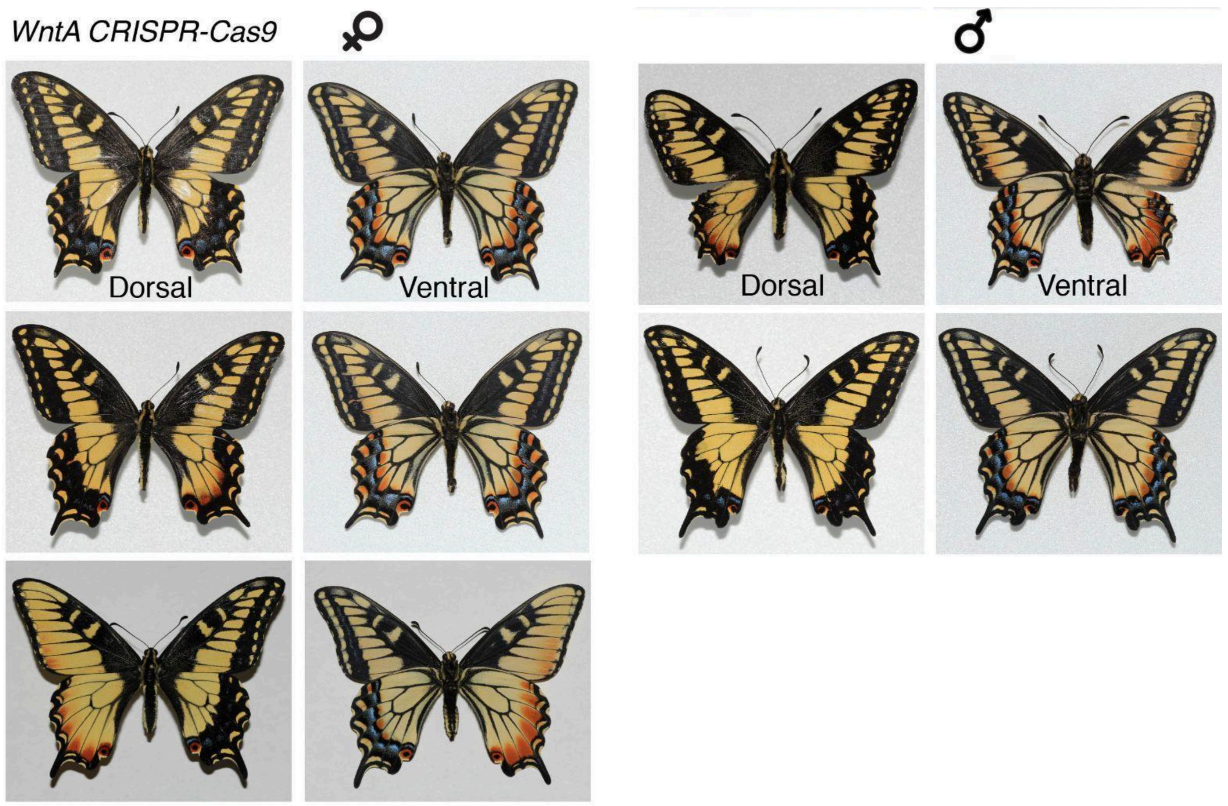


**Figure S12.** Effects of *WntA* CRISPR/Cas9 on *Papilio zelicaon*.

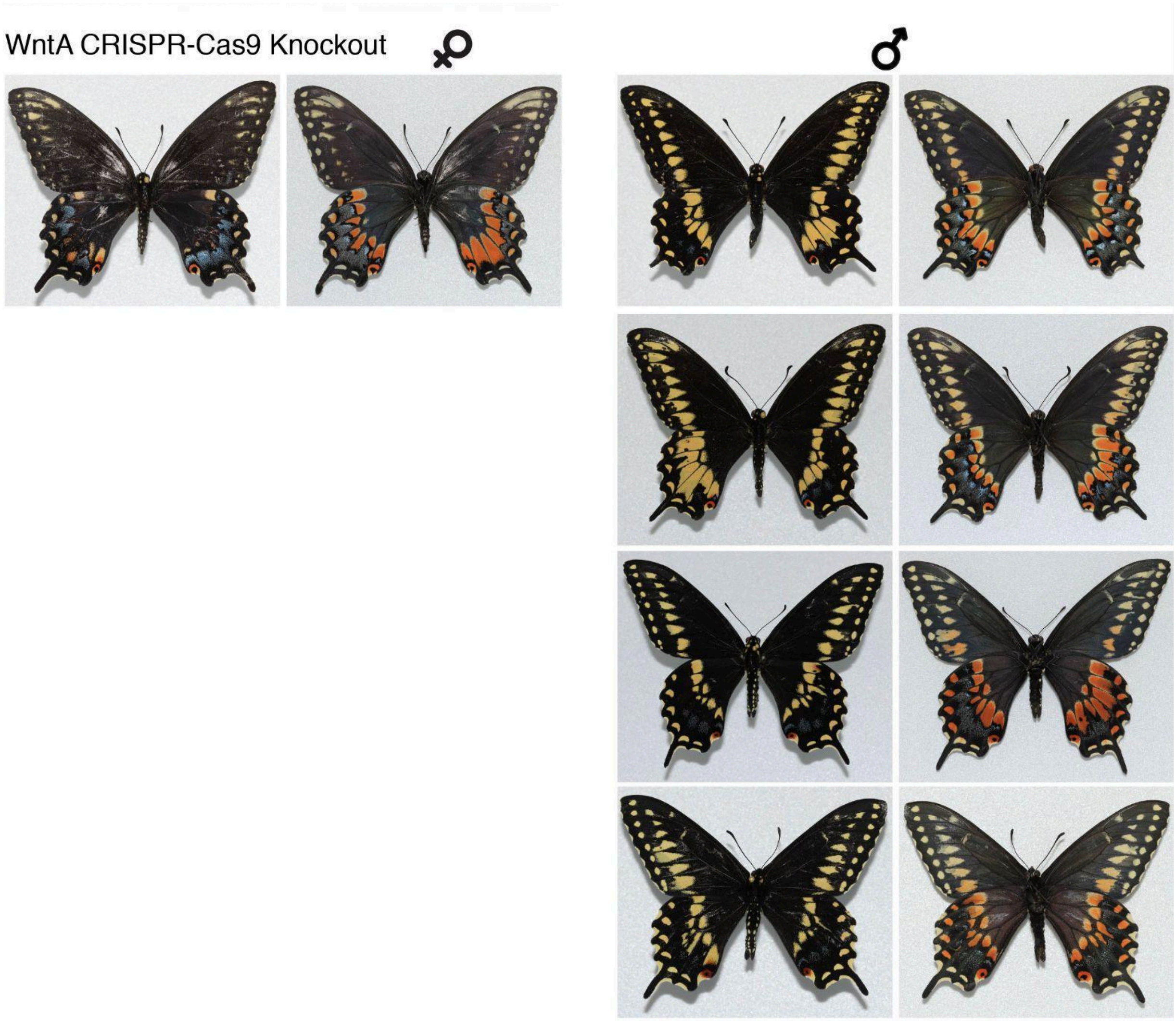


**Figure S13.** Effects of *WntA* CRISPR/Cas9 on *Papilio polyxenes*.

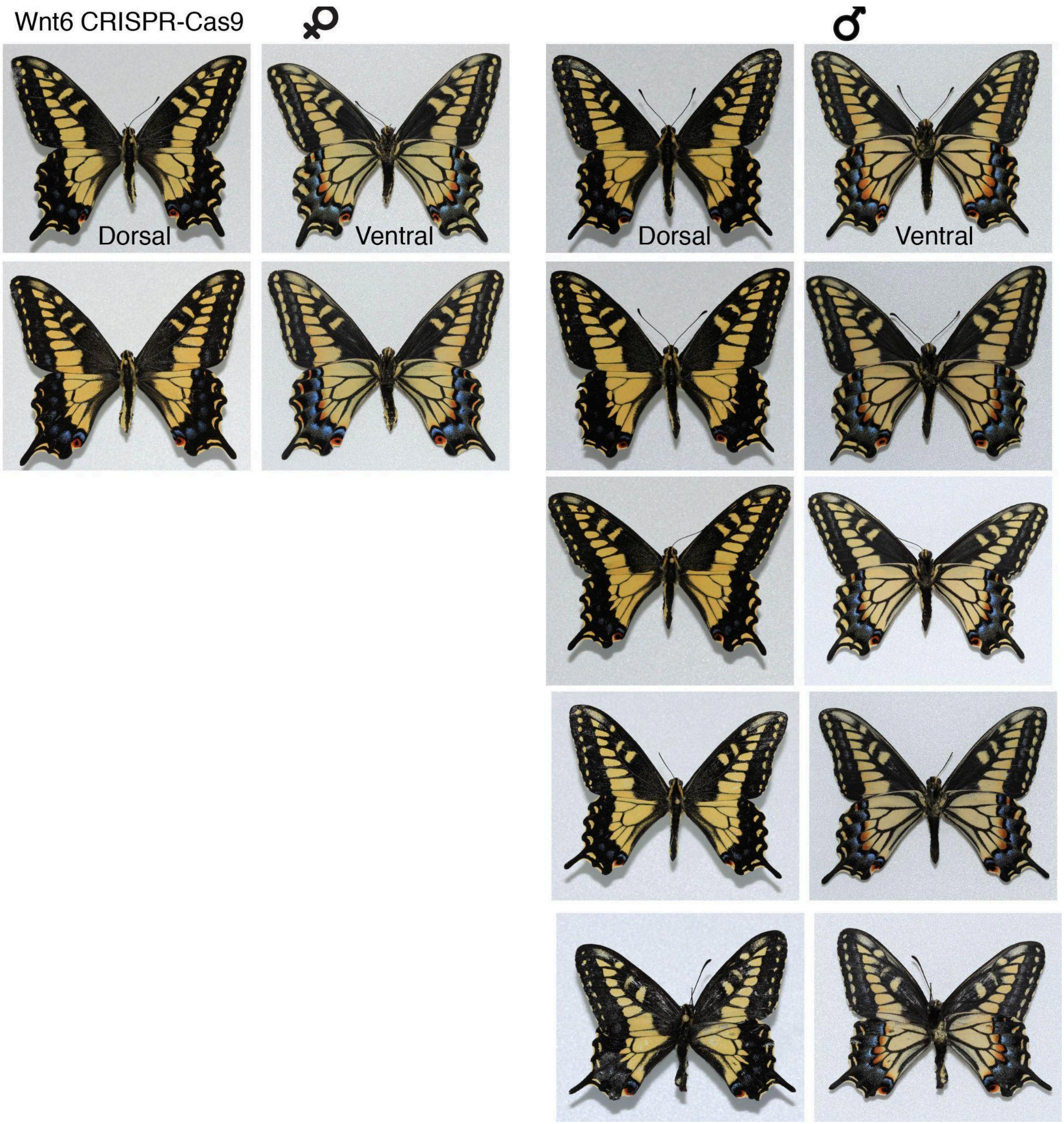


**Figure S14.** Effects of *Wnt6* CRISPR/Cas9 on *Papilio zelicaon*.

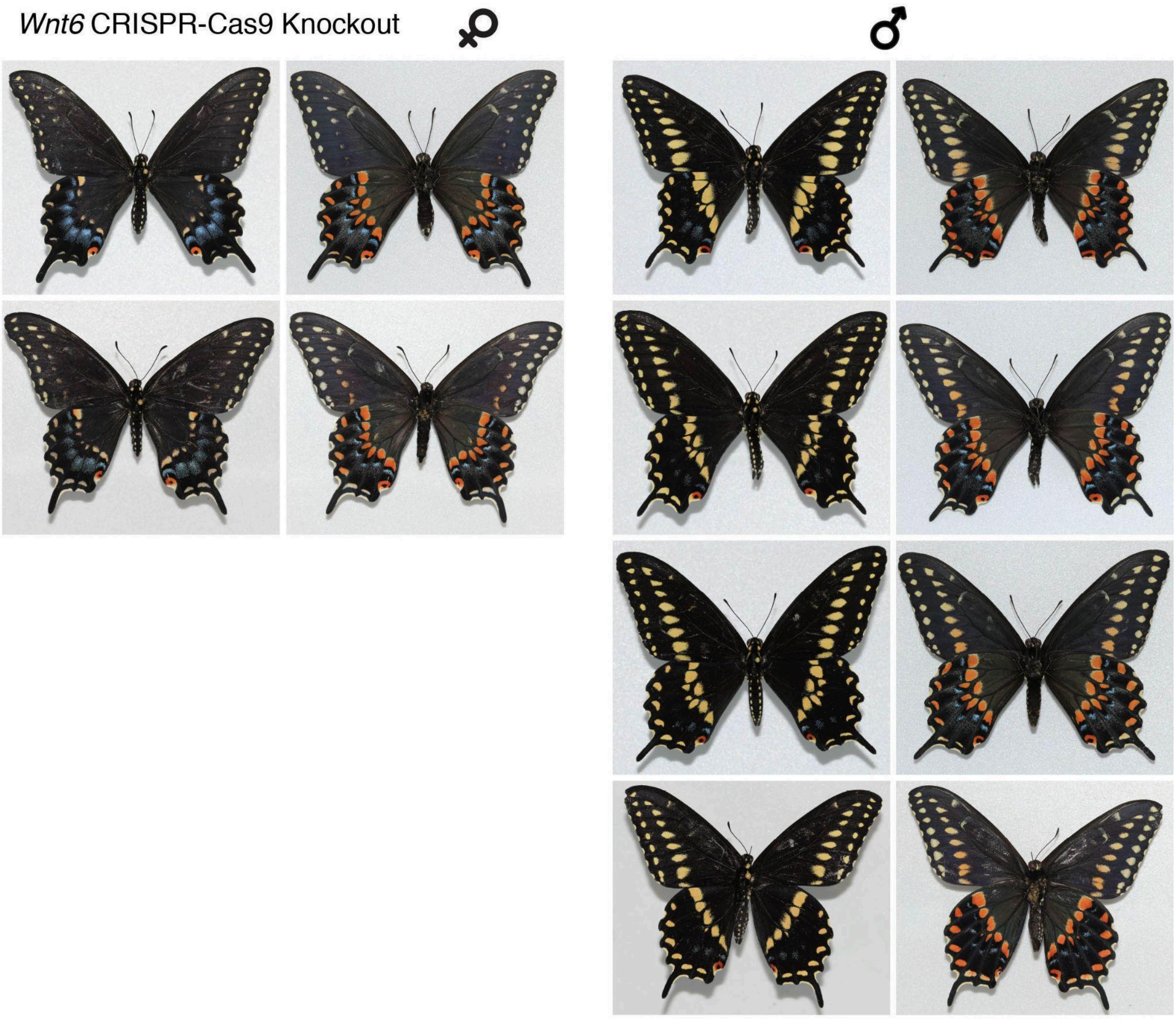


**Figure S15.** Effects of *Wnt6* CRISPR/Cas9 on *Papilio polyxenes*.
